# AsCRISPR: a web server for allele-specific sgRNA design in precision medicine

**DOI:** 10.1101/672634

**Authors:** Guihu Zhao, Jinchen Li, Yu Tang

## Abstract

Allele-specific genomic targeting by CRISPR provides a point of entry for personalized gene therapy of dominantly inherited diseases, by selectively disrupting the mutant alleles or disease-causing single nucleotide polymorphisms (SNPs), ideally while leaving normal alleles intact. Moreover, the allele-specific engineering has been increasingly exploited not only in treating inherited diseases and mutation-driven cancers, but also in other important fields such as genome imprinting, haploinsufficiency, genome loci imaging and immunocompatible manipulations. Despite the tremendous utilities of allele-specific targeting by CRISPR, very few bioinformatic tools have been implemented for the allele-specific purpose. We thus developed **AsCRISPR** (**<u>A</u>**llele-**<u>s</u>**pecific **CRISPR**), a web tool to aid the design of guide RNA (gRNA) sequences that can discriminate between alleles. It provides users with limited bioinformatics skills to analyze both their own identified variants and heterozygous SNPs deposited in the dbSNP database. Multiple CRISPR nucleases and their engineered variants including newly-developed Cas12b and CasX are included for users’ choice. Meanwhile, **AsCRISPR** evaluates the on-target efficiencies, specificities and potential off-targets of gRNA candidates, and also displays the allele-specific restriction enzyme sites that might be disrupted upon successful genome edits. In addition, **AsCRISPR** analyzed with dominant single nucleotide variants (SNVs) retrieved from ClinVar and OMIM databases, and generated a ***Dominant Database*** of candidate discriminating gRNAs that may specifically target the alternative allele for each dominant SNV site. A ***Validated Database*** was also established, which manually curated the discriminating gRNAs that were experimentally validated in the mounting literatures. **AsCRISPR** is freely available at http://www.genemed.tech/ascrispr.

## 1. Introduction

Inherited diseases are caused by various types of mutations, insertions/deletions (indels), large genomic structural variations, as well as pathogenic single nucleotide polymorphisms (SNPs) that are critical for personalized medicine. Of those, dominant inherited diseases present a special challenge for researchers to conduct gene therapies. Those patients inherited only one mutated allele and one normal allele on pairs of chromosomes. The treatment strategy typically involves an allele-specific manipulation by silencing or ablating the pathogenic alleles while exerting no aberrant effects on the wildtype ones. Previously, numerous studies used allele-specific short interference RNAs (siRNAs) to selectively suppress dominant mutant allele, and produced immense therapeutic benefits [1, 2]. Until recent years, allele-specific CRISPR genome editing has emerged to be a promising means to treat human genetic disorders, and has also been increasingly implemented in other versatile applications such as genome imprinting, haploinsufficiency, genome loci imaging and immunocompatible manipulations [3]. The CRISPR system provides a highly specific genome editing that is capable of discriminating disease-causing alleles from wildtype ones, whenever the genetic variants (1) **In PAM**: generate unique protospacer adjacent motifs (PAMs); or (2) **Near PAM**: located within the spacer region, especially the seed region of short guide RNAs (sgRNAs) [3, 4].

So far, allele-specific CRISPR has been increasingly employed in treating various diseases such as retinitis pigmentosa [5–8], corneal dystrophy [9], dominant progressive hearing loss [10] and multiple cancers [11–13], as well as genome imprinting diseases [14]. And it was also used to alleviate haploinsufficiency by allele-specific CRISPR activation of wildtype alleles [15], and even was designed for manipulating human leukocyte antigen (HLA) locus [16]. More excitingly, this strategy has been recently utilized to selectively inactivate mutant Huntington (mHTT), taking advantage of novel PAMs created by SNPs flanking the HTT locus [17, 18]. Overall, allele-specific CRISPR is now believed to be a promising personalized strategy for treating genetic diseases.

However, it is always labor intensive and time consuming to figure out appropriate guide sequences that may discriminate between two alleles [9, 17]. Currently, most web servers only design sgRNAs from the reference genomes, without allele discriminations. Thus we developed **AsCRISPR** (**<u>A</u>**llele-**<u>s</u>**pecific **CRISPR**), a web server to aid the design of sgRNAs, for allele-specific genome engineering. It has incorporated multiple CRISPR nucleases and can process flexibly with either user-identified variants or heterozygous SNPs. **AsCRISPR** facilitates the selection of optimal discriminating sgRNAs, with the output of multiple on-target scores and elaborated off-target candidates. Importantly, we also developed a database of dominant SNVs, for the first time, that are targetable by the CRISPR system in an allele-specific manner. Meanwhile, we are in the ongoing process of collecting experimentally validated allele-specific sgRNAs reported in the mounting literature for treating diseases.

### 2. Implementation

**AsCRISPR** was developed using PHP and Perl on a Linux platform with an Apache web server. The front and back separation model was used; the front end is based on the Vue + Element, and the back end is based on the Laravel, a PHP web framework.

Single-base mutations, short indels, and SNP IDs are the formats for input (**Figure 1**). The SNP information was downloaded from dbSNP v150 database (https://www.ncbi.nlm.nih.gov/SNP) and stored in MySQL database. To optimize the SNP query performance, an index on SNP table was added. Sequence can be extracted from the .2bit file (hg19/GRCh37, hg38/GRCh38, or mm10/GRCm38) with the twoBitToFa command base on the SNP information (chromosome, start genomic position and end genomic position, reference allele and alternate allele). **AsCRISPR** displayed the SNP sites located at both flanking nucleotides of a query SNP ID, which was implemented using D3.

**Figure 1.**
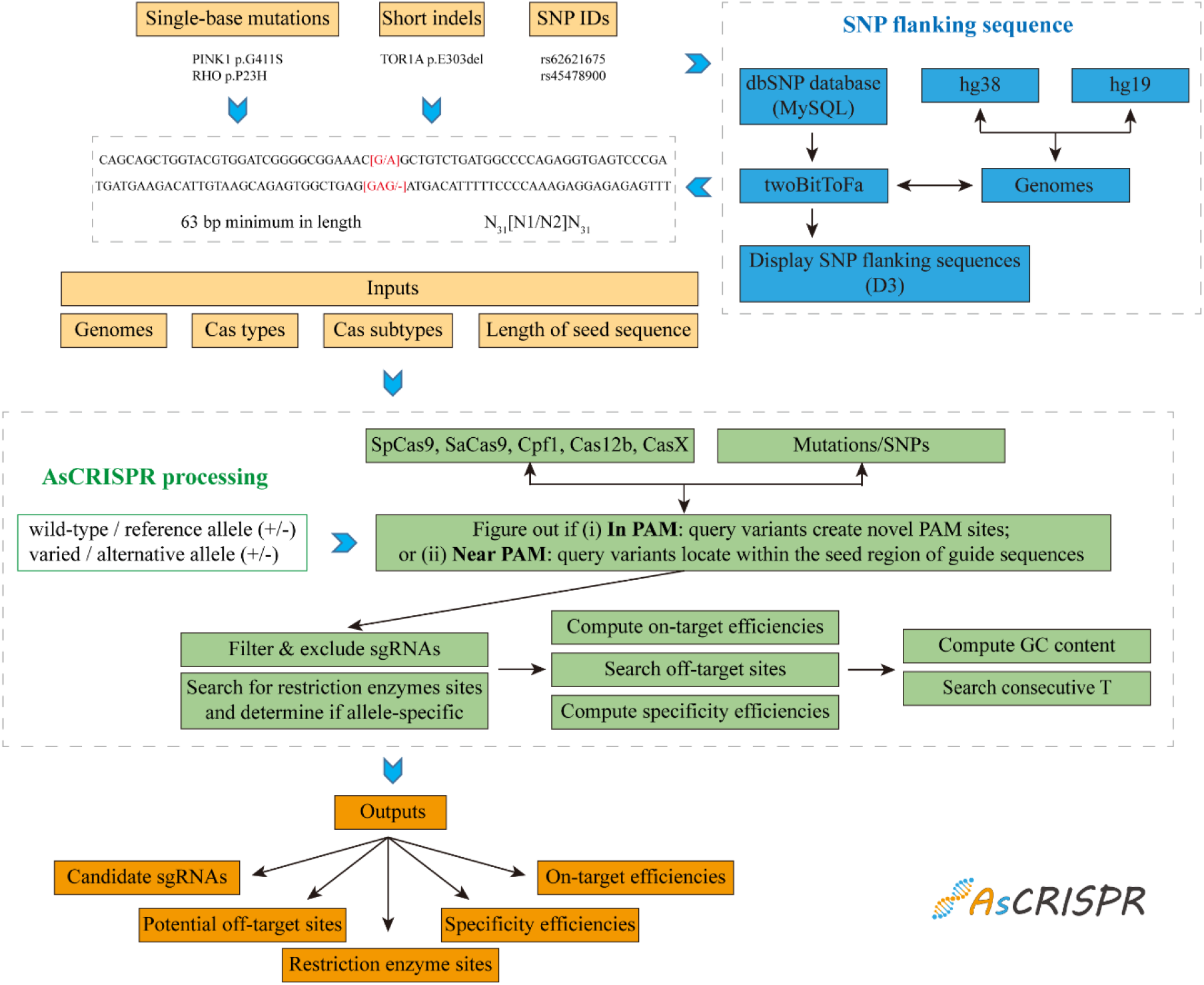
The workflow of AsCRISPR.

In principle, **AsCRISPR** proceeds to figure out if (i) **In PAM**: query variants give rise to novel PAMs, which confers stringent allele-specific targeting, or (ii) **Near PAM**: query variants locate within the seed region of guide sequences, which may abolish the Cas cleavage (**Figure 1 and Supplementary Figure 1A**). **AsCRISPR** then outputs the candidate guide sequences, after performing the stringent search and filter. For example, those guide sequences with novel PAMs generated by variants that constitute an ambiguous genetic code (such as R and Y in the CjCas9:NNNNRYAC), will be excluded.

Scripts from CRISPOR (https://github.com/maximilianh/crisporPaper) were then integrated into **AsCRISPR** to assess sgRNA properties and scores. **AsCRISPR** also searched for possible sites recognized by restriction enzymes deposited in our database. In addition, guide sequences were further analyzed and reminded as “Not recommended” if (i) the GC contents are beyond 20%~80%; or (ii) containing four or more consecutive T, which might terminate the U6 or U3 promoter-drived transcription.

The dataset for ***Dominant Database*** was sourced from the ClinVar database [19]. We chose ‘pathogenic’ and ‘pathogenic/likely pathogenic’ variants deposited in the ClinVar database after June 6^th^ 2020. Autosomal dominant information of variants was extracted from Online Mendelian Inheritance in Man (OMIM, V20190524) [20].

## 3. Results

**AsCRISPR** helps to design sgRNAs based on four major types of Cas nucleases including the commonly used Cas9, Cpf1 and also recently reported Cas12b [21, 22] and CasX [23], each type of which contains its variant subtypes with distinct PAM sites and seed lengths (**Table 1**). This allows the users to freely choose the optimal combination of Cas protein and sgRNA to meet their own needs.

**Table 1.**
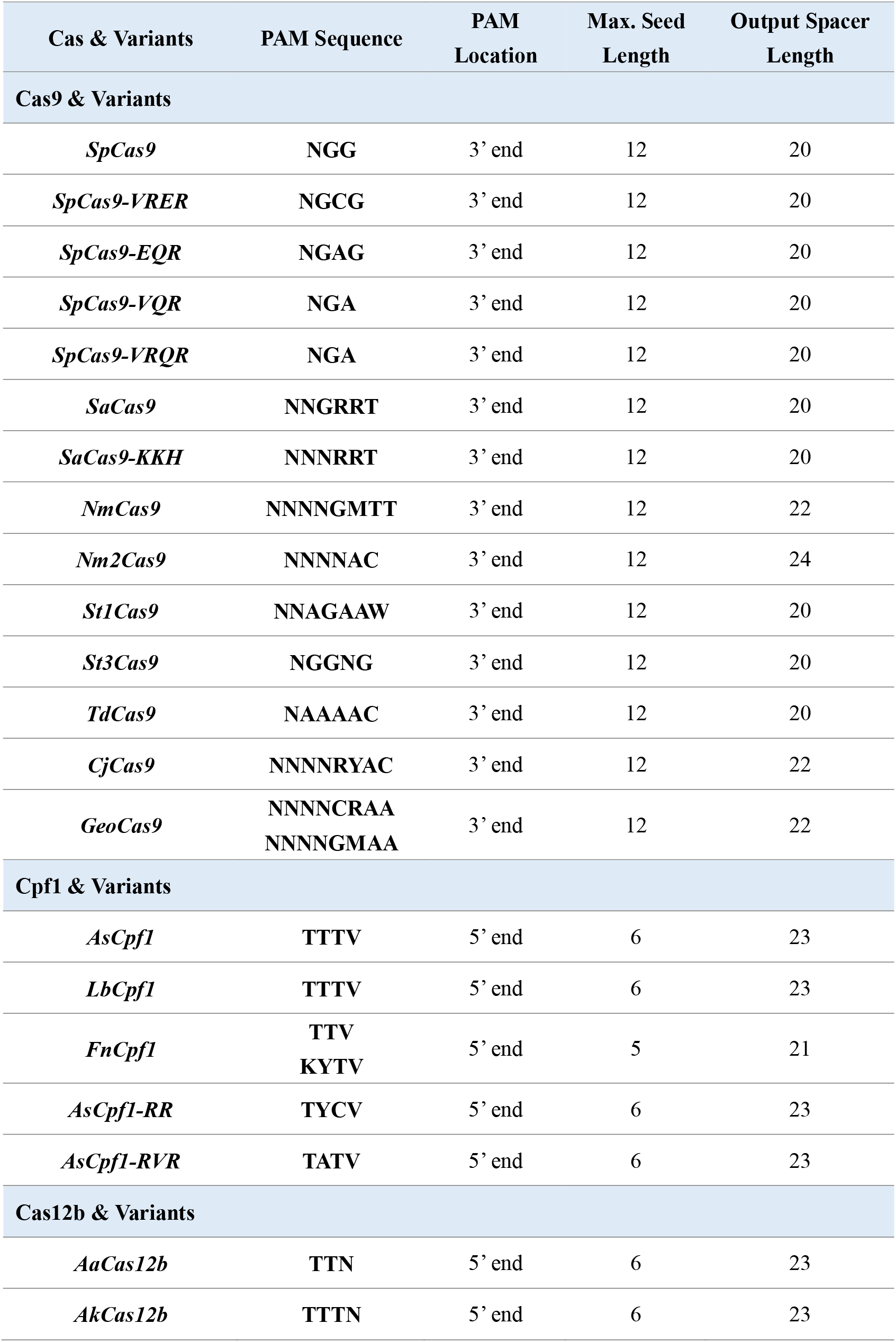
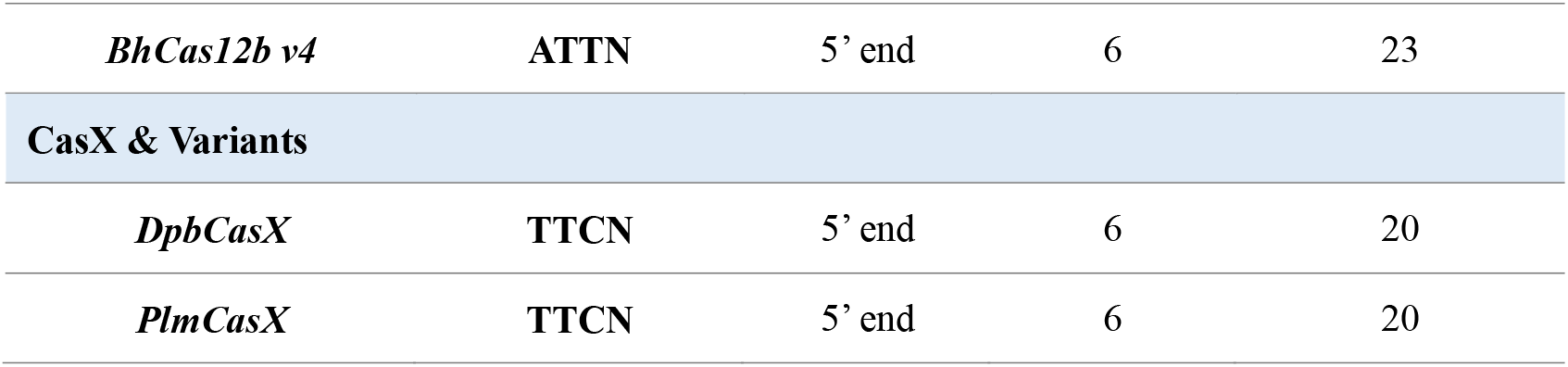
Cas nucleases and their variants used in AsCRISPR.

### 3.1 Input Format

The inputs for **AsCRISPR** could be DNA sequences harboring single-base mutations or short indels, and also simply SNP IDs deposited in the dbSNP database. All inputs will be finally processed as the format of N_31_[N1/N2]N_31_, in which N1/N2 denotes the sequence in the wildtype/reference and mutated/alternative allele, respectively. Thus, it means that the input sequence requires 63 bp minimum in length, with at least 31 bp flanking the variant site, to be processed for the output of a complete list of candidate discriminating sgRNAs (**Supplementary Figure 1B, C**). Notably, when users are in query with an SNP ID, **AsCRISPR** will also display other SNP sites located at both flanking 31 nucleotides (**Supplementary Figure 1D**), which provides extra variation information and would be of great value for designing personalized genome targeting.

### 3.2 Candidate Guides

**AsCRISPR** provides downloadable results with candidate sgRNAs that target only one allele (**Figure 2**). For better visualization, **AsCRISPR** ranks all guide sequences first by listing pairs with the same PAM sequence back-to-back. Furthermore, **AsCRISPR** evaluates their on-target efficiencies, specificity efficiencies and potential off-targets throughout the genome, taking advantage of the CRISPOR’s scoring system [24]. Specifically, the on-target efficiencies were calculated with multiple reported algorithms and were normalized to 0-1. For SpCas9, efficiency scores were predicted according to Xu et al., 2015 [25]; Doench et al., 2016 [26]; Moreno-Mateos et al., 2015 [27]; and Listgarden et al., 2018 [28], respectively. For SaCas9, efficiency scores were predicted according to Najm et al., 2018 [29]. And for Cpf1, Cas12b and CasX, efficiency scores were predicted according to Kim et al., 2018 [30].

**Figure 2.**
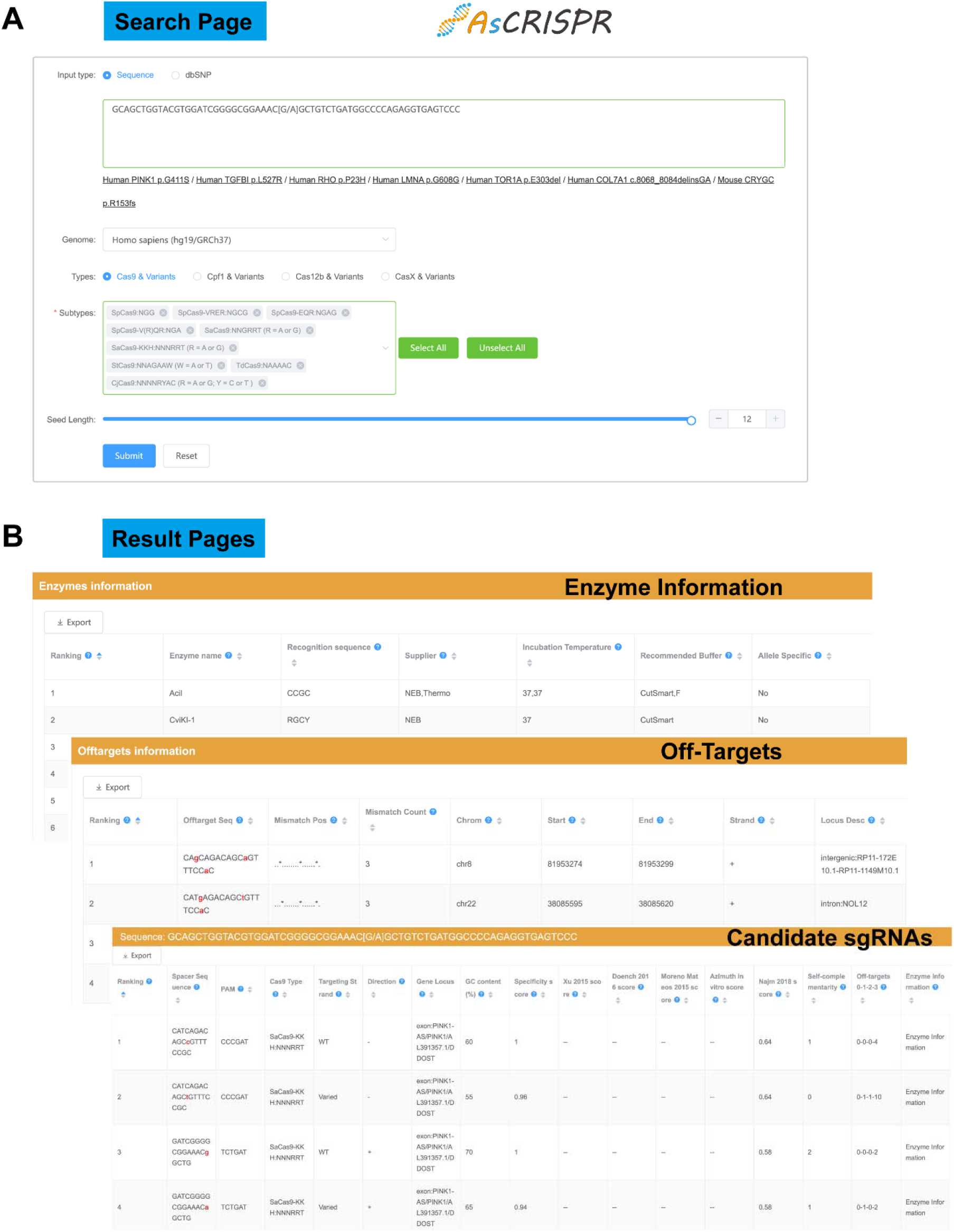
Snapshot of the graphic user interface of AsCRISPR. **(A)** Search page. The **AsCRISPR** homepage allows two types of input (DNA sequence or SNP IDs) with default parameters optimized for novice users. **(B)** Result pages including the output of guide sequences, off-targets information and restriction enzyme information.

### 3.3 Off-targets

The potential off-target sequences throughout the genome are rigorously searched by 3-base mismatches maximum (**Figure 2**). **AsCRISPR** lists the number of off-targets for each guide sequence with 0, 1, 2 or 3 mismatches (0-1-2-3). Clicking on the (0-1-2-3) will reveal more information about the off-target information in the downstream data sheet, including the locations (exon, intron or intergenic region), sequence mismatches and so forth. Users can freely re-rank the off-targets by locations. The specificity score measures the uniqueness of a guide sequence in the genome. The higher the specificity score, the lower are off-target effects. Specificity scores were calculated based on Hsu scores [31] and CFD scores [26]. For Cpf1, Cas12b and CasX, no off-target ranking algorithms were available in the literatures so far, instead we just applied Hsu and CFD scores to their off-targets.

### 3.4 Restriction Sites

**AsCRISPR** also searches for possible sites recognized by restriction enzymes along the spacer sequences (**Figure 2**), which might be disrupted after gene targeting, and further determines whether those candidate enzymes are also allele-specific. This provides an important tool for the characterization and screening of targeted single colonies by restriction fragment length polymorphism (RFLP).

### 3.5 Exemplary Running

We have listed several typical sequences on the website for exemplary running. For example, heterozygous *PINK1 p.G411S* is one of the ideal mutations for allele-specific targeting, which was previously demonstrated to increase the risk of Parkinson’s disease via a dominant-negative mechanism [32]. In the Cas9 mode, **AsCRISPR** outputs 11 discriminating sgRNAs in combination with 3 subtypes of Cas9 including SpCas9, SpCas9-V(R)QR and SaCas9-KKH (**Supplementary Table 1**). One of those sgRNA exploits a novel PAM (5’-CgG-3’) created by the mutation, and another 5 pairs of sgRNAs containing the mutation point within the seed region selectively target either wildtype allele or mutated allele (**Supplementary Table 1**). Therefore, by using Cas9, totally 5 candidate sgRNAs might be specific to the mutated *PINK1 p.G411S* allele, which are ready for the users’ experimental evaluations (**Supplementary Table 1**). Besides, we also listed other exemplary mutations including the single mutations (*TGFBI p.L527R; RHO p.P23H; LMNA p.G608G*), 3-base delete mutations (*TOR1A p.E303del*) and short indel mutations (*COL7A1 c.8068_8084delinsGA*).

Similarly, for heterozygous SNPs, **AsCRISPR** processes with the input SNP numbers and translates them into DNA sequences (63 bp) after retrieving the genomic database. As an example, we used **AsCRISPR** to analyze one of the SNPs, rs62621675:[C>G], only with Cas9, and successfully obtained 24 discriminating sgRNAs in combination with 6 subtypes of Cas9 including SpCas9-V(R)QR, SpCas9-EQR, SpCas9-VRER, CjCas9, SaCas9-KKH and Nme2Cas9 (**Supplementary Table 1**). Four of those sgRNA exploits novel PAMs (5’-AgA-3’; 5’-AgAG-3’; 5’-AGAGACAc-3’; 5’-AGACAc-3’) created by the SNP, and another 10 pairs of sgRNAs containing the variant point within the seed sequence (**Supplementary Table 1**).

Users can freely re-rank the candidate sgRNAs by Cas types, on-target efficiencies, specificity scores, off-targets properties, and others. We recommend to select the sgRNAs with novel PAMs, since they contribute to the most stringent discrimination. For a more detailed demonstration, users can also find a **AsCRISPR** tutorial on the website, which can be read online or downloaded as a pdf document (**Supplementary File 1**).

### 3.6 Databases

To strengthen the purpose of targeting dominant diseases, **AsCRISPR** collected dominant SNV sites from ClinVar and OMIM databases for allele-specific analysis, and generated a ***Dominant Database*** depositing candidate targetable sgRNAs, which may specifically target the alternative allele for each dominant SNV site. Notably, we observed that the ClinVar and OMIM databases are not correctly annotated, in several cases, with the dominant nature of diseases. Thus several clearly dominant diseases/SNV sites might be filtered out from the ***Dominant Database***. Nevertheless, in the current version of **AsCRISPR**, we collected 102212 records of dominant SNV sites (with 12938 SNV sites were pathogenic or likely-pathogenic), related to 1833 different diseases. **AsCRISPR** analyzed the dominant SNV sites and found that around 36.96% (38.25% for pathogenic dominant SNVs) are targetable by allele-specific CRISPR (**Figure 3A**). Among the four major types of Cas nucleases, Cas9 is the most frequently employed, probably due to its multiple subtypes (**Figure 3B**). Further analysis showed that four Cas9 subtypes including SpCas9, SpCas9-V(R)QR, SaCas9-KKH and Nme2Cas9 are obviously more robust in targeting SNV sites, which is consistent with their simpler nature of PAM sequences (**Figure 3C**). Generally, the alternative allele of each dominant SNV site can be targeted by an average of 14.15 discriminating sgRNAs, among those 2.84 sgRNAs are **In PAM** (create a novel PAM site) and 11.57 sgRNAs are **Near PAM** (in spacer seed region) (**Figure 3D**). Interestingly, the major types of Cas nucleases that contribute to the stringent **In PAM** targeting are SpCas9, SpCas9-V(R)QR, SaCas9-KKH and Nme2Cas9 (**Figure 3E**). Basically, the targeting features are similar between dominant SNVs and pathogenic dominant SNVs (**Figure 3**).

**Figure 3.**
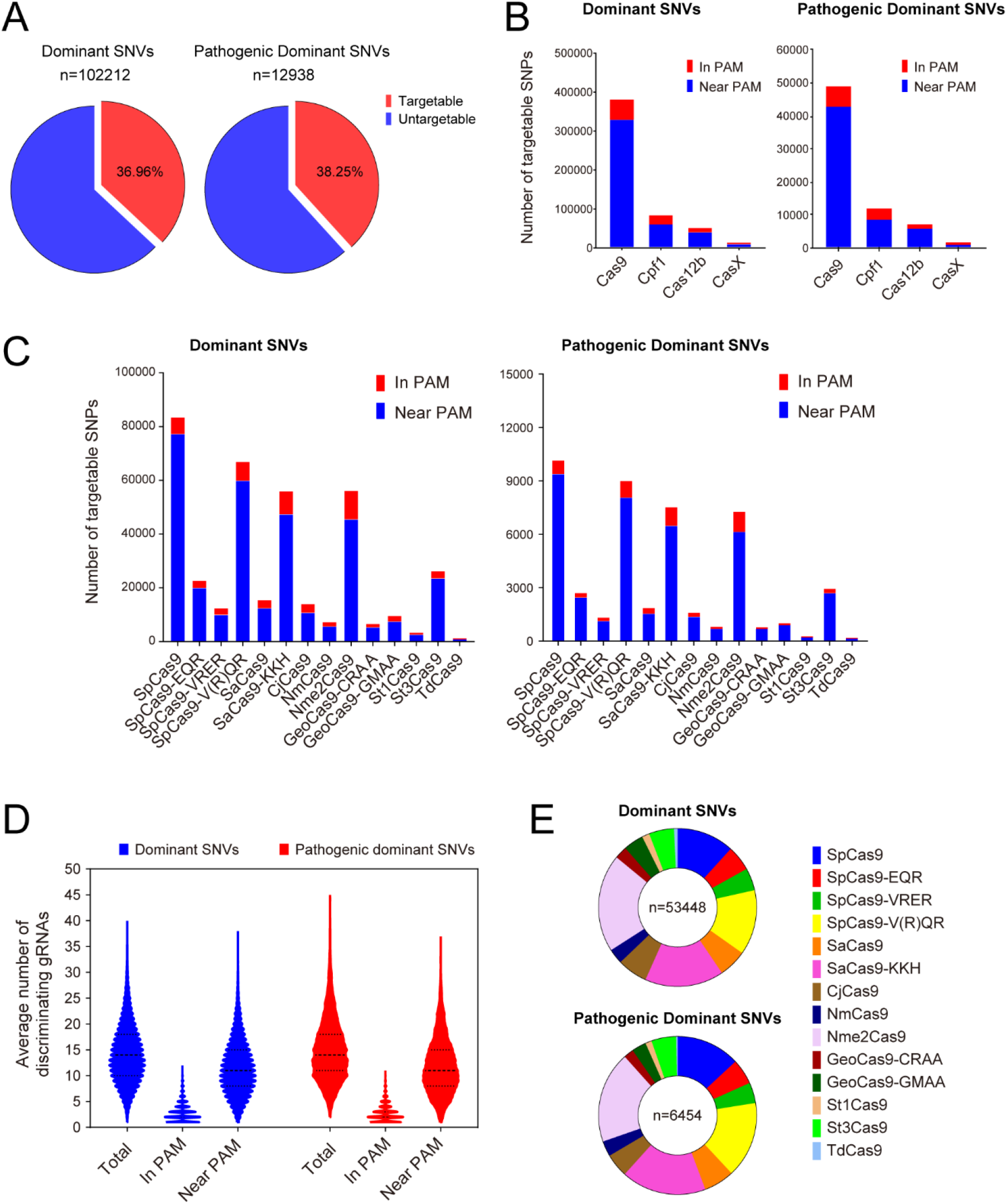
Statistics of *Dominant Database*. **(A)** Ratios of targetable dominant SNVs and pathogenic dominant SNVs. **(B, C)** Targetable SNVs by different Cas types and subtypes. **(D)** Average number of discriminating gRNAs for each SNV site. **(E)** In PAM gRNAs by different Cas types and subtypes.

Considering the predictive nature of ***Dominant Database***, we also manually curated the experimentally validated sgRNAs that were reported in the mounting literatures for treating diseases in an allele-specific manner, and thus established a ***Validated Database***. This database is expected to be expanded rapidly along with the advancements in this area.

## 4. Future Developments

So far, **AsCRISPR** has integrated the genomes of Homo sapiens (hg19/GRCh37), Homo sapiens (hg38/GRCh38) and Mus musculus (mm10/GRCm38). We are planning to upload more genomes for analysis in the near future, to expand its allele-specific utilities. However, users can still input their identified sequences or DNA sequences from other species, the current version of **AsCRISPR** would then output the discriminating sgRNAs, but without displaying the information of gene locus, scores and off-targets.

Increasing types of Cas nucleases such as ScCas9 [33], xCas [34], SauriCas9 [35] and recently developed Cas12j [36], as well as those targeting on RNAs such as CasRx [37] and Cas13a [38, 39] (already reported with allele-specific purpose), will be added for expanded utilities.

Our understanding of on- and off-target sgRNA efficiencies is evolving rapidly. Although the on-target efficiencies in **AsCRISPR** were calculated with multiple reported algorithms, the scoring algorithms have been continuously improved. Cas12b and CasX may have their special efficiency scoring algorithms that are different from those of Cas9 and Cpf1, however, to our best knowledge, there are still lack of published studies working on it. We will thus incorporate the convincing scoring algorithms, which predict either on- or off-target efficiencies, into **AsCRISPR** once they became available.

**AsCRISPR** will be timely updated along with the emerging exploitation of Cas toolbox. Moreover, the ***Dominant Database*** and ***Validated Database*** resources will be updated three times per year. We also welcome users to send us their experimentally validated discriminating sgRNAs for our update. All in all, we welcome any constructive feedback from users for improving our web server.

## 5. Conclusion and Discussion

### 5.1 Comparison with similar servers

We have thus developed **AsCRISPR**, which is an easy-to-use and streamlined web tool for designing potential discriminating sgRNAs between alleles to facilitate the CRISPR-based personalized therapy and other versatile applications. Particularly, we incorporated two recently reported types of Cas nucleases, Cas12b and CasX, which show to be promising for genetic engineering due to their smaller size and higher specificity.

As we just finished the **AsCRISPR** implementation, another software termed AlleleAnalyzer was published, aiming to identify optimized pairs of personalized and allele-specific sgRNAs [40]. AlleleAnalyzer also leverages patterns of shared genetic variation across thousands of publicly available genomes to design sgRNA pairs that will have the greatest utility in large populations [40]. However, the difference is that **AsCRISPR** as a web tool can process either user identified sequences or SNP numbers, which is more likely demand-driven for research studies and clinical therapeutics. Moreover, **AsCRISPR** only outputs single sgRNAs instead of pairs of sgRNAs, although the users may also freely use **AsCRISPR** to manually design another sgRNA to make an sgRNA pair. Note that numerous non-coding RNAs or regulatory elements are widespread, dual-sgRNA excision of a large DNA fragment might bring about extra risks for disease treatment. Thus, **AsCRISPR** may possess extra allele-specific utilities and add to the bioinformatic repositories for allele-specific genomic editings.

Another two web servers, SNP-CRISPR and CrisPam, were also recently developed, converging with our web tool, **AsCRISPR**, but at different publication status, to share the same notion of the allele-specific designing of sgRNAs. Basically, SNP-CRISPR designed the sgRNAs based on public SNP database yet with only two types of PAMs [41]. Another web server, CrisPam, only analyzed the variants that produce novel PAMs (**In PAM**) from pathogenic and likely-pathogenic SNPs in humans [42]. Interestingly, they found that 84% of the total SNPs checked can create novel PAMs and the average number of PAMs generated by a SNP is 6.97 [42]. In contrast, analysis results with our database turns out that an average of 2.84 sgRNAs are **In PAM**. The difference comes from multiple aspects. We analyzed only dominant SNVs from ClinVar and OMIM databases, which underlies potential differences. Yet, the major difference is the type of Cas proteins used for analysis. In CrisPam, multiple Cas with simpler PAMs such as xCas were involved. Regarding this, the results from our database are still underestimated. Nevertheless, more tool Cas proteins with expanded PAM sequences are in needy to achieve the full potential of allele-specific genome targeting.

The brief properties of different servers were summarized in **Table 2**. **AsCRISPR** works to be the most comprehensive web tool for allele-specific purpose, endowed with most of Cas nucleases and PAM sites, selectable seed lengths, importantly the quality control information of candidate sgRNAs such as multiple on-target scores, elaborated off-target sequences and RFLP sites, as well as curated databases of targetable dominant SNVs and experimentally validated discriminating sgRNAs.

**Table 2.**
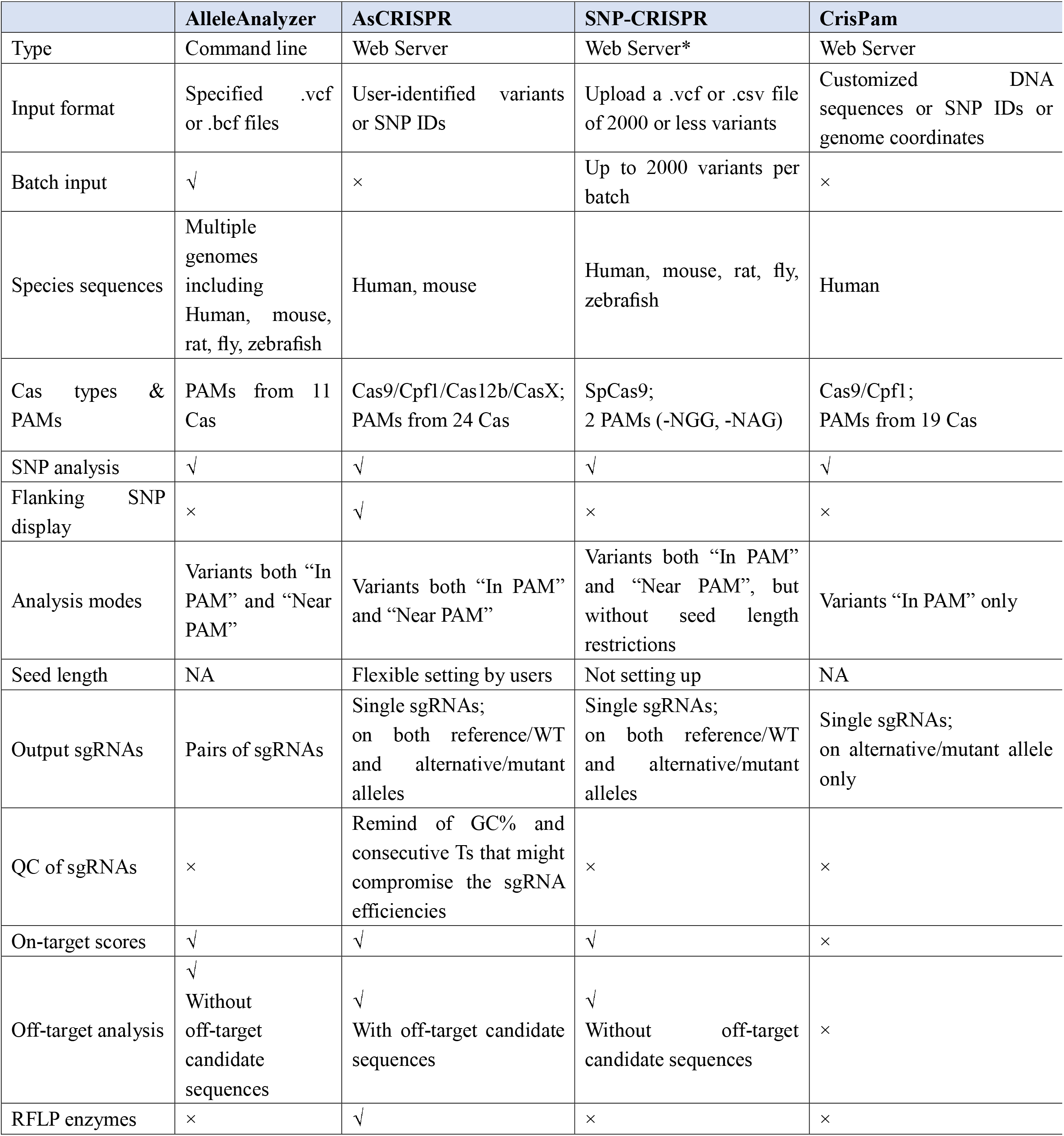
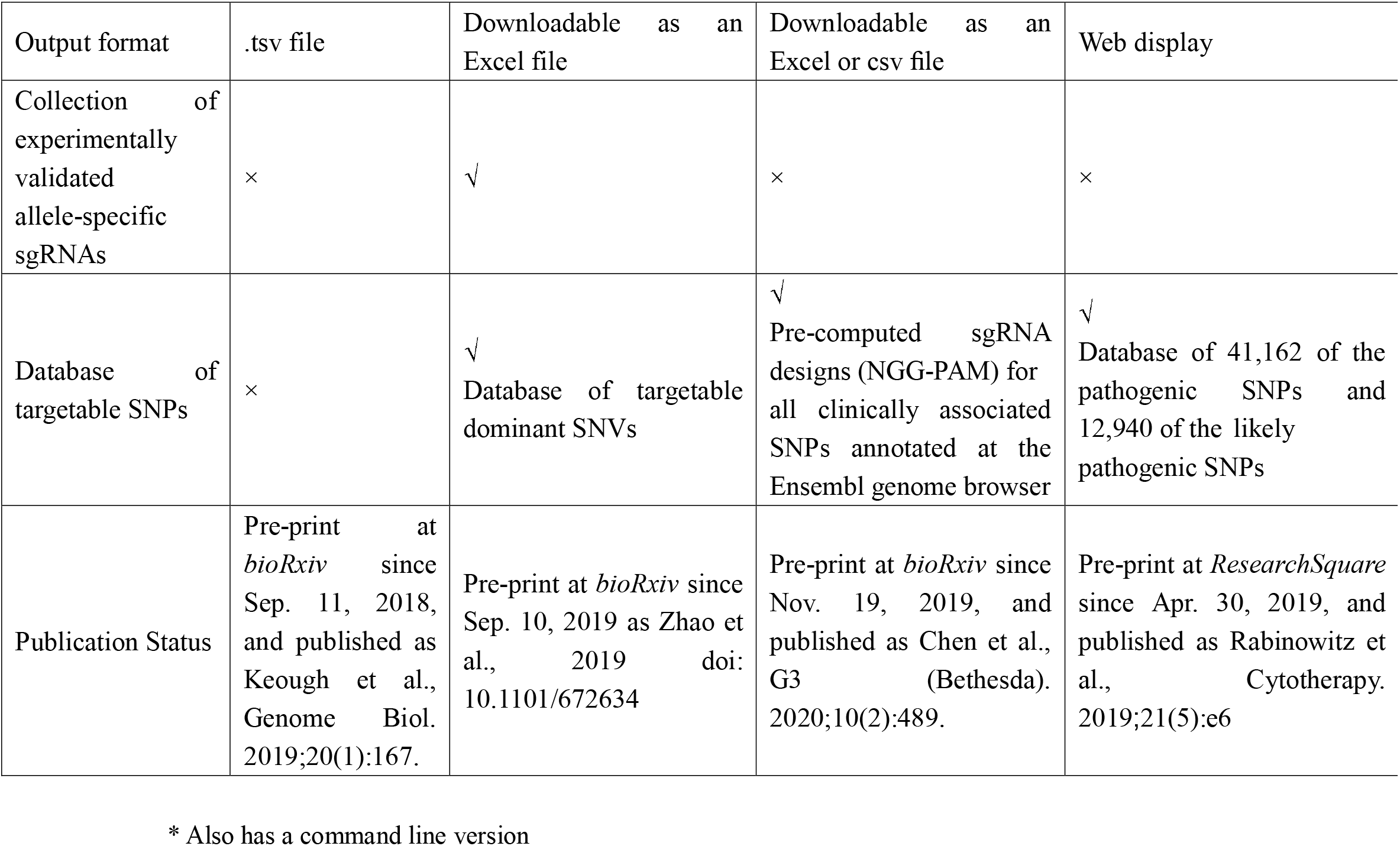
Comparison of tools associated with allele-specific genome targeting.

### 5.2 Population vs. Personalized Therapy

It has been a long time that, interestingly, people have been avoiding genetic variants when designing sgRNAs for therapeutic genome editing in large populations. Previous studies have performed comprehensive analysis on the Exome Aggregation Consortium (ExAC) and 1000 Genomes Project (1000GP) data sets, and determined that genetic variants could negatively impact sgRNA efficiency, as well as both on- and off-target specificity at therapeutically implicated loci [43, 44]. Thus, for the CRISPR-based therapy in large patient populations, genetic variations should be considered in the design and evaluation of sgRNAs to minimize the risk of treatment failure and/or adverse outcomes. To address that, people thus endeavor to identify universal/platinum sgRNAs located in the low-variation regions, with the help of, for example, the ExAC or gnomAD browser, to maximize their population efficacy [43, 45].

Although the genetic variations would be a challenge for platinum sgRNA design, it provides a promising entry for designing allele-specific or personalized sgRNAs in treating individual patients. Deciphering genetic variations helps to seek common platinum sgRNAs for the treatment in large populations, whereas the tools including **AsCRISPR** go towards the opposite direction that exploits the discrimination abilities of heterozygous genetic variants to facilitate the design of allele-specific targets for individuals.

The rapid advancements of genome sequencing technologies have accelerated the research of variants and their physiological functions or effects in the era of precision medicine [46–49]. Specifically, with the expanding toolkits of the CRISPR system, emerging studies has been performed regarding the variants linking with the pathogenesis of human diseases. Apart from treating genetic diseases, allele-specific CRISPR genome editing has also been increasingly used in other research areas such as genome imprinting, haploinsufficiency, spatiotemporal loci imaging and immunocompatible manipulations, and, for sure, would be exploited for increasing applications in the near future. Therefore, **AsCRISPR** would be right on track along with the rapidly progressing research on genetic variants.

## Supporting information

Supplementary Figure 1_AsCRISPR

Supplementary File 1_AsCRISPR_Tutorial Page

Supplementary Table 1 Exemplary Running

## Authors’ Contributions

YT conceived of, designed, and directed the study. GZ wrote the scripts and implemented the website. YT and GZ wrote and revised the manuscript. JL helped with the data retrieving and analysis. All authors read and approved the final manuscript.

## Acknowledgments

This work was supported by grants from the National Natural Sciences Foundation of China [81801200 to Y. T.]; talents startup funds of Xiangya Hospital [2209090550057 to Y. T.]; and Hunan Provincial Natural Science Foundation of China [2019JJ40476 to Y. T., 2019JJ50974 to G. Z.].

## References

1. Monga, I., et al., ASPsiRNA: A Resource of ASP-siRNAs Having Therapeutic Potential for Human Genetic Disorders and Algorithm for Prediction of Their Inhibitory Efficacy. G3 (Bethesda), 2017. 7(9): p. 2931–2943.

2. Hohjoh, H., Disease-causing allele-specific silencing by RNA interference. Pharmaceuticals (Basel), 2013. 6(4): p. 522–35.

3. Wu, J., B. Tang, and Y. Tang, Allele-specific genome targeting in the development of precision medicine. Theranostics, 2020. 10(7): p. 3118–3137.

4. Cong, L., et al., Multiplex genome engineering using CRISPR/Cas systems. Science, 2013. 339(6121): p. 819–23.

5. Burnight, E.R., et al., Using CRISPR-Cas9 to Generate Gene-Corrected Autologous iPSCs for the Treatment of Inherited Retinal Degeneration. Mol Ther, 2017. 25(9): p. 1999–2013.

6. Giannelli, S.G., et al., Cas9/sgRNA selective targeting of the P23H Rhodopsin mutant allele for treating retinitis pigmentosa by intravitreal AAV9.PHP.B-based delivery. Hum Mol Genet, 2018. 27(5): p. 761–779.

7. Li, P., et al., Allele-Specific CRISPR-Cas9 Genome Editing of the Single-Base P23H Mutation for Rhodopsin-Associated Dominant Retinitis Pigmentosa. CRISPR J, 2018. 1: p. 55–64.

8. Bakondi, B., et al., In Vivo CRISPR/Cas9 Gene Editing Corrects Retinal Dystrophy in the S334ter-3 Rat Model of Autosomal Dominant Retinitis Pigmentosa. Mol Ther, 2016. 24(3): p. 556–63.

9. Christie, K.A., et al., Towards personalised allele-specific CRISPR gene editing to treat autosomal dominant disorders. Sci Rep, 2017. 7(1): p. 16174.

10. Gyorgy, B., et al., Allele-specific gene editing prevents deafness in a model of dominant progressive hearing loss. Nat Med, 2019. 25(7): p. 1123–1130.

11. Gao, Q., et al., Selective targeting of the oncogenic KRAS G12S mutant allele by CRISPR/Cas9 induces efficient tumor regression. Theranostics, 2020. 10(11): p. 5137–5153.

12. Lentsch, E., et al., CRISPR/Cas9-Mediated Knock-Out of Kras(G12D) Mutated Pancreatic Cancer Cell Lines. Int J Mol Sci, 2019. 20(22).

13. Koo, T., et al., Selective disruption of an oncogenic mutant allele by CRISPR/Cas9 induces efficient tumor regression. Nucleic Acids Res, 2017. 45(13): p. 7897–7908.

14. Bashtrykov, P., G. Kungulovski, and A. Jeltsch, Correction of aberrant imprinting by allele-specific epigenome editing. Clin Pharmacol Ther, 2016. 99(5): p. 482–4.

15. Matharu, N., et al., CRISPR-mediated activation of a promoter or enhancer rescues obesity caused by haploinsufficiency. Science, 2019. 363(6424).

16. Xu, H., et al., Targeted Disruption of HLA Genes via CRISPR-Cas9 Generates iPSCs with Enhanced Immune Compatibility. Cell Stem Cell, 2019. 24(4): p. 566–578 e7.

17. Monteys, A.M., et al., CRISPR/Cas9 Editing of the Mutant Huntingtin Allele In Vitro and In Vivo. Mol Ther, 2017. 25(1): p. 12–23.

18. Shin, J.W., et al., Permanent inactivation of Huntington’s disease mutation by personalized allele-specific CRISPR/Cas9. Hum Mol Genet, 2016. 25(20): p. 4566–4576.

19. Landrum, M.J., et al., ClinVar: improving access to variant interpretations and supporting evidence. Nucleic Acids Res, 2018. 46(D1): p. D1062–D1067.

20. Amberger, J.S., et al., OMIM.org: Online Mendelian Inheritance in Man (OMIM(R)), an online catalog of human genes and genetic disorders. Nucleic Acids Res, 2015. 43(Database issue): p. D789–98.

21. Strecker, J., et al., Engineering of CRISPR-Cas12b for human genome editing. Nat Commun, 2019. 10(1): p. 212.

22. Teng, F., et al., Repurposing CRISPR-Cas12b for mammalian genome engineering. Cell Discov, 2018. 4: p. 63.

23. Liu, J.J., et al., CasX enzymes comprise a distinct family of RNA-guided genome editors. Nature, 2019. 566(7743): p. 218–223.

24. Haeussler, M., et al., Evaluation of off-target and on-target scoring algorithms and integration into the guide RNA selection tool CRISPOR. Genome Biol, 2016. 17(1): p. 148.

25. Xu, H., et al., Sequence determinants of improved CRISPR sgRNA design. Genome Res, 2015. 25(8): p. 1147–57.

26. Doench, J.G., et al., Optimized sgRNA design to maximize activity and minimize off-target effects of CRISPR-Cas9. Nat Biotechnol, 2016. 34(2): p. 184–191.

27. Moreno-Mateos, M.A., et al., CRISPRscan: designing highly efficient sgRNAs for CRISPR-Cas9 targeting in vivo. Nat Methods, 2015. 12(10): p. 982–8.

28. Listgarten, J., et al., Prediction of off-target activities for the end-to-end design of CRISPR guide RNAs. Nat Biomed Eng, 2018. 2(1): p. 38–47.

29. Najm, F.J., et al., Orthologous CRISPR-Cas9 enzymes for combinatorial genetic screens. Nat Biotechnol, 2018. 36(2): p. 179–189.

30. Kim, H.K., et al., Deep learning improves prediction of CRISPR-Cpf1 guide RNA activity. Nat Biotechnol, 2018. 36(3): p. 239–241.

31. Hsu, P.D., et al., DNA targeting specificity of RNA-guided Cas9 nucleases. Nat Biotechnol, 2013. 31(9): p. 827–32.

32. Puschmann, A., et al., Heterozygous PINK1 p.G411S increases risk of Parkinson’s disease via a dominant-negative mechanism. Brain, 2017. 140(1): p. 98–117.

33. Chatterjee, P., N. Jakimo, and J.M. Jacobson, Minimal PAM specificity of a highly similar SpCas9 ortholog. Sci Adv, 2018. 4(10): p. eaau0766.

34. Hu, J.H., et al., Evolved Cas9 variants with broad PAM compatibility and high DNA specificity. Nature, 2018. 556(7699): p. 57–63.

35. Hu, Z., et al., A compact Cas9 ortholog from Staphylococcus Auricularis (SauriCas9) expands the DNA targeting scope. PLoS Biol, 2020. 18(3): p. e3000686.

36. Pausch, P., et al., CRISPR-CasPhi from huge phages is a hypercompact genome editor. Science, 2020. 369(6501): p. 333–337.

37. Konermann, S., et al., Transcriptome Engineering with RNA-Targeting Type VI-D CRISPR Effectors. Cell, 2018. 173(3): p. 665–676 e14.

38. Abudayyeh, O.O., et al., C2c2 is a single-component programmable RNA-guided RNA-targeting CRISPR effector. Science, 2016. 353(6299): p. aaf5573.

39. Zhao, X., et al., A CRISPR-Cas13a system for efficient and specific therapeutic targeting of mutant KRAS for pancreatic cancer treatment. Cancer Lett, 2018. 431: p. 171–181.

40. Keough, K.C., et al., AlleleAnalyzer: a tool for personalized and allele-specific sgRNA design. Genome Biol, 2019. 20(1): p. 167.

41. Chen, C.L., et al., SNP-CRISPR: A Web Tool for SNP-Specific Genome Editing. G3 (Bethesda), 2020. 10(2): p. 489–494.

42. R., R., D. R., and O. D., CrisPam: SNP-derived PAM analysis web tool and human pathogenic SNPs database for CRISPR allele-specific targeting. Cytotherapy, 2019. 21(5).

43. Scott, D.A. and F. Zhang, Implications of human genetic variation in CRISPR-based therapeutic genome editing. Nat Med, 2017. 23(9): p. 1095–1101.

44. Lessard, S., et al., Human genetic variation alters CRISPR-Cas9 on- and off-targeting specificity at therapeutically implicated loci. Proc Natl Acad Sci U S A, 2017. 114(52): p. E11257–E11266.

45. Karczewski, K.J., et al., The mutational constraint spectrum quantified from variation in 141,456 humans. Nature, 2020. 581(7809): p. 434–443.

46. Xu, X., et al., The A/G allele of eIF3a rs3740556 predicts platinum-based chemotherapy resistance in lung cancer patients. Lung Cancer, 2013. 79(1): p. 65–72.

47. Guo, C., et al., Pharmacogenomics signature: A novel strategy on the individual differences in drug response. Cancer Lett, 2018. 420: p. 190–194.

48. Wang, Y., et al., Genetic polymorphisms of lncRNA-p53 regulatory network genes are associated with concurrent chemoradiotherapy toxicities and efficacy in nasopharyngeal carcinoma patients. Sci Rep, 2017. 7(1): p. 8320.

49. Wang, Y., et al., The association of transporter genes polymorphisms and lung cancer chemotherapy response. PLoS One, 2014. 9(3): p. e91967.

