## Supplementary Figure 1_AsCRISPR for "AsCRISPR: a web server for allele-specific sgRNA design in precision medicine"

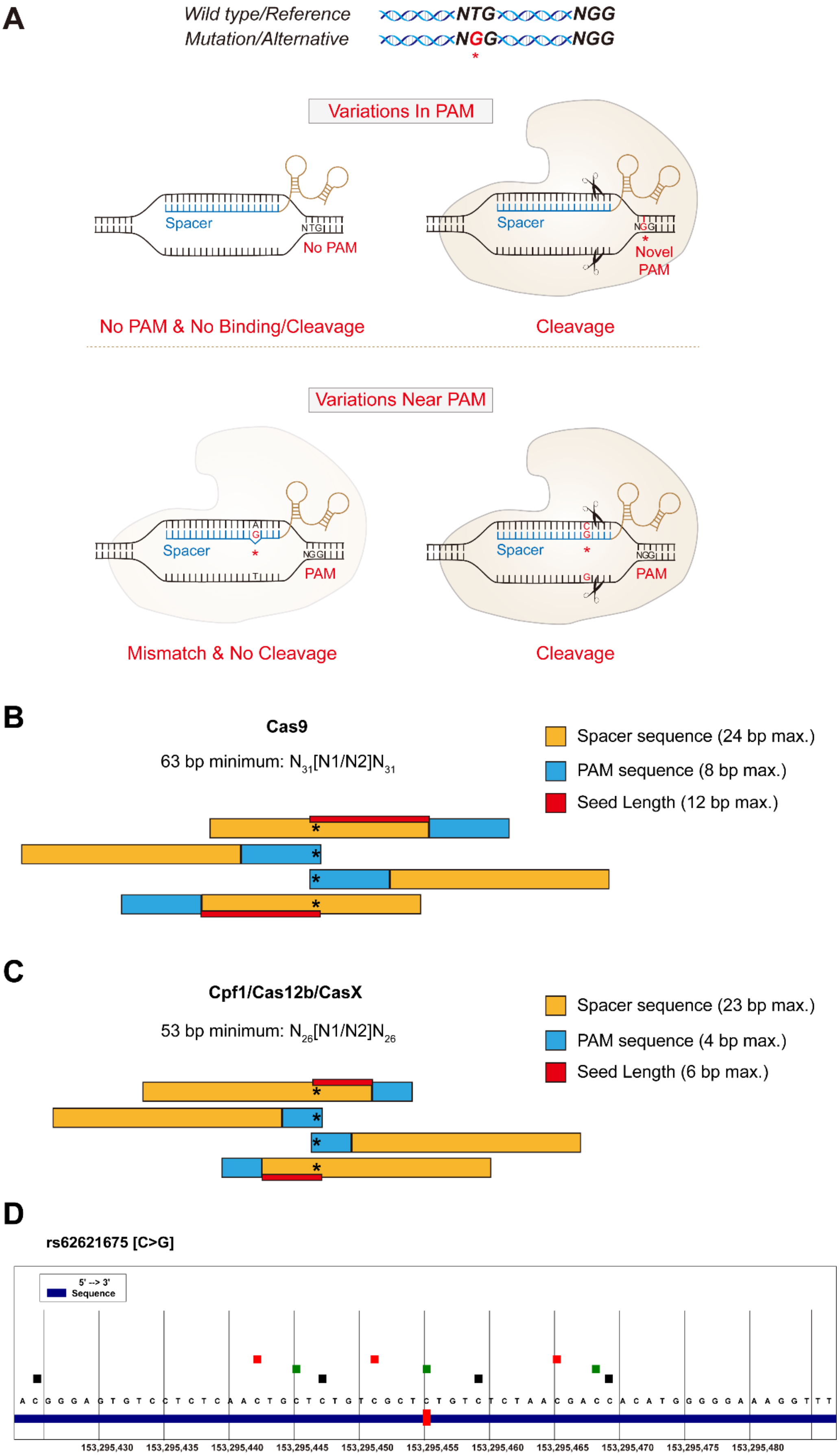
A
Wild type/Reference
NTG
NGG
Mutation/Alternative
NGG
NGG
*
Variations In PAM
Spacer
NTG
No PAM
Spacer
NGG
*
Novel PAM
No PAM & No Binding/Cleavage
Cleavage
Variations Near PAM
Spacer
A
G
*
T
NGG
PAM
Spacer
C
G
*
G
NGG
PAM
Mismatch & No Cleavage
Cleavage
B
Cas9
63 bp minimum: N31[N1/N2]N31
Spacer sequence (24 bp max.)
PAM sequence (8 bp max.)
Seed Length (12 bp max.)
C
Cpf1/Cas12b/CasX
53 bp minimum: N26[N1/N2]N26
Spacer sequence (23 bp max.)
PAM sequence (4 bp max.)
Seed Length (6 bp max.)
D
rs62621675 [C>G]
5' --> 3'
Sequence
A C G G G A G T G T C C T C T C A A C T G C T C T G T C G C T C T G T C T C T A A C G A C C A C A T G G G G G A A A G G T T T
153,295,430
153,295,435
153,295,440
153,295,445
153,295,450
153,295,455
153,295,460
153,295,465
153,295,470
153,295,475
153,295,480

**Supplementary Figure 1. Allele-specific models and input of query sequences. (A)** The "**In PAM**" model and "**Near PAM**" model for allele-specific genomic targeting. The "**In PAM**" model confers the most stringent allele-specific cleavages, since the binding of Cas with its target DNA is diminished. The "**Near PAM**" model exploits the discrimination ability of sgRNAs, with mutations or SNPs that locate within the spacer region, particularly the seed region. The variant is denoted by a red asterisk. *Adapted from Wu et al., Allele-specific genome targeting in the development of precision medicine. Theranostics, 2020. 10(7): p. 3118-3137.* **(B, C)** Demonstration of query DNA sequences for Cas9: 63 bp minimum **(B)** and Cpf1/Cas12b/CasX: 53 bp minimum **(C)**. To maximize users' demands, the length of input spacer sequence for Cas9 was set as 24 bp for query. The input DNA sequence for AsCRISPR basically requires minimum 63 bp in length, with at least 31 bp flanking the mutation/variation site (*), to completely showcase the possible candidate sgRNAs. The format of input DNA sequences would be: $N_{31}$[N1/N2]$N_{31}$, in which N1 denotes the sequences in the wildtype allele and N2 denotes the sequences in the mutated or varied allele. **(D)** In query with an SNP ID, AsCRISPR will display other SNP sites located at both flanking 31 nucleotides.
