## Supplementary File 1_AsCRISPR_Tutorial Page for "AsCRISPR: a web server for allele-specific sgRNA design in precision medicine"

### About AsCRISPR

**AsCRISPR** (**A**llele-**s**pecific **CRISPR**) is a web tool for designing guide sequences for allele-specific CRISPR genome engineering. It is developed to aid the design of guide sequences during the allele-specific gene engineering of dominant inherited diseases, and can be further exploited in many other versatile applications, such as treating genome imprinting and haploinsufficiency, spatiotemporal loci imaging and immunocompatible manipulations. For dominantly inherited diseases, patients inherit one pathogenic allele causing a disease phenotype, especially in a dominant-negative manner, and one normal allele as well. Treatment strategies typically involve silencing the mutated allele, whereas expression of the wild-type (WT) allele remains unaffected and is sufficient for normal functions to prevent the onset of the disease. Allele-specific genome editing by CRISPR is a promising means to treat those diseases by selectively targeting the mutated genes or disease-causing single nucleotide polymorphisms (SNPs). Notably, in the era of personalized medicine, naturally occurring SNPs that alter the protospacer adjacent motifs (PAMs) or spacer sequences recognized by CRISPR nucleases would also provide a point of entry for patient-specific gene therapy.

Basically, mutations/SNPs-derived novel PAMs can confer stringent allele-specific targeting, and also guide sequences with mutations/SNPs located especially within their seed sequence can also distinguish between two alleles. **AsCRISPR** can process with either user-identified sequences harboring single-base mutations and short insertion-deletion mutations (indels), or heterozygous SNPs deposited in the dbSNP database. Multiple CRISPR nucleases including SpCas9, SaCas9, Cpf1, Cas12b, CasX and their engineered variants are included for users' choice. **AsCRISPR** can then return the candidate guide sequences that may selectively target one of the alleles, and further evaluate their on-target efficiencies, specificity efficiencies and potential off-targets throughout the genome, and it can also display the restriction enzyme sites that might be disrupted after gene targeting, which is of great use for clonal screening.

Moreover, **AsCRISPR** analyzed with dominant single nucleotide variants (SNVs) retrieved from ClinVar and OMIM and provided a ***Dominant Database*** of candidate targetable sgRNAs, which may specifically target the alternative allele for each dominant SNV site. Besides, **AsCRISPR** also established a ***Validated Database*** that deposited all experimentally validated allele-specific sgRNAs for treating genetic diseases.

### How to use AsCRISPR?

#### Input

Input type: The inputs can be either DNA sequences or SNP IDs deposited in the dbSNP database. The input DNA sequence requires minimum 63 bp in length, with at least 31 bp flanking the mutation/variation site. The format of input DNA sequence would be: $N_{31}$[N1/N2]$N_{31}$, in which N1 denotes the sequences in the wildtype allele and N2 denotes the sequences in the mutated or varied allele.

For example:

- Human RHO p.P23H

TTCTCCAATGCGACGGGTGTGGTACGCAGCC[C/A]CTTCGAGTACCCACAGTACTACCTGGCTGAG

- Human TOR1A p.ΔE302/303

TGATGAAGACATTGTAAGCAGAGTGGCTGAG[GAG/-]ATGACATTTTTCCCCAAAGAGGAGAGAGTTT

Alternatively, users can input an SNP ID including “rs”: rs62621675 or rs28937568, and click “Get Sequences”. **AsCRISPR** then displays the SNP site and retrieve its flanking sequences, in the format of $N_{31}$[N1/N2]$N_{31}$ for users’ choice and further analysis. For example:

- rs62621675

ACGGGAGTGTCCTCTCAACTGCTCTGTCGCT[C/G]TGTCTCTAACGACCACATGGGGGAAAGGTTT

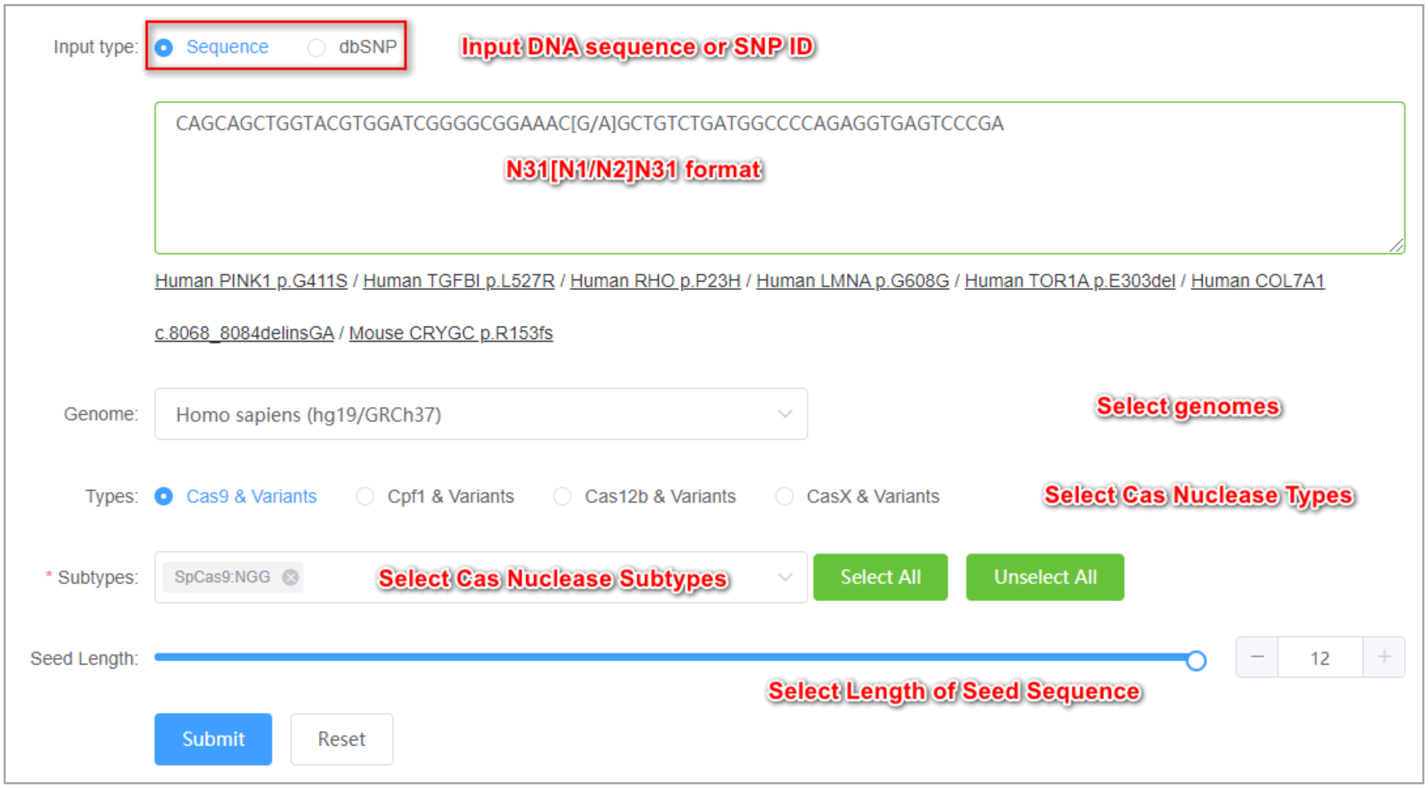

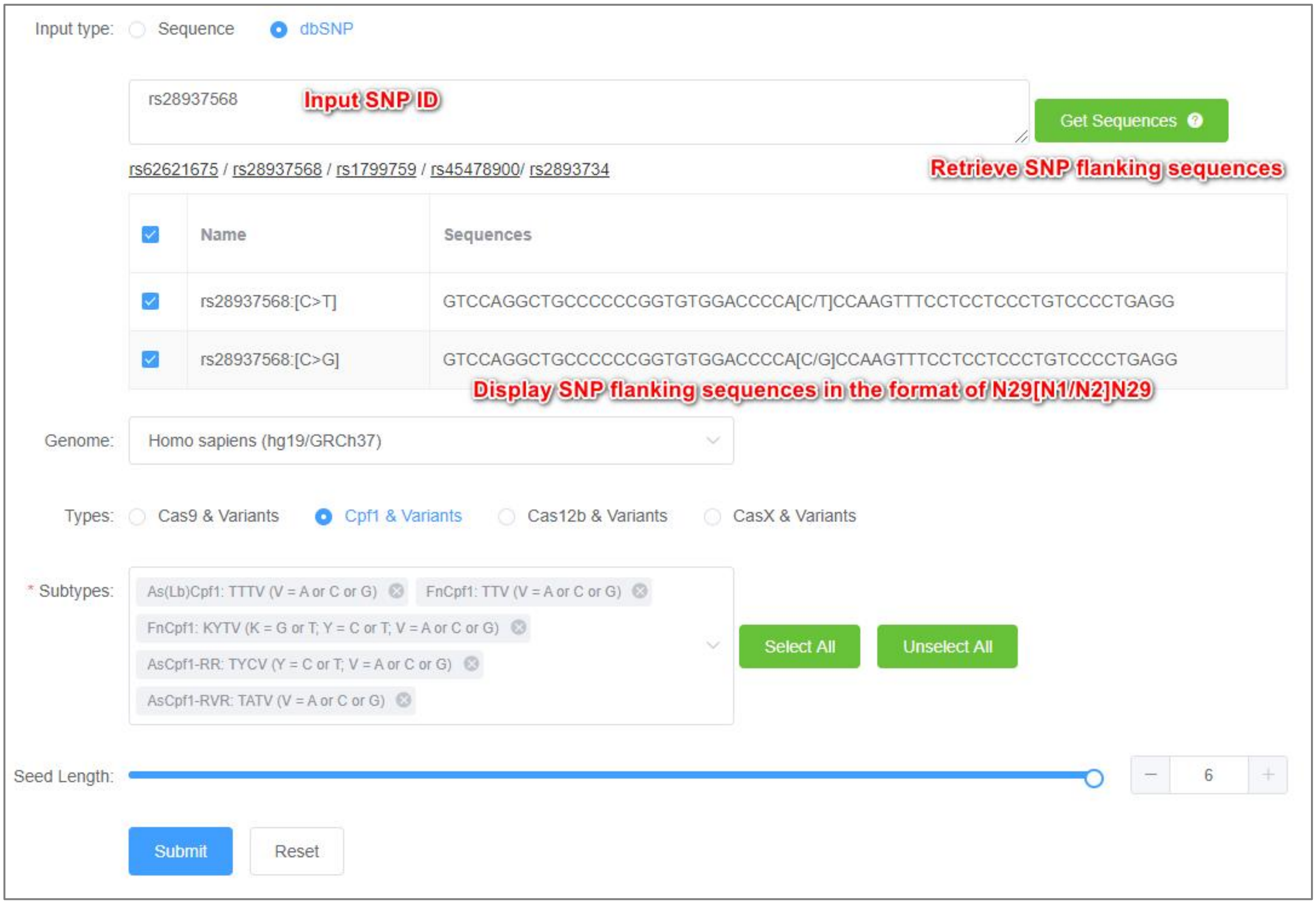


Moreover, when users are in a query with an SNP ID, **AsCRISPR** will also display the SNP sites located at each flanking 31 nucleotides, which would be informative for personalized gene targeting.

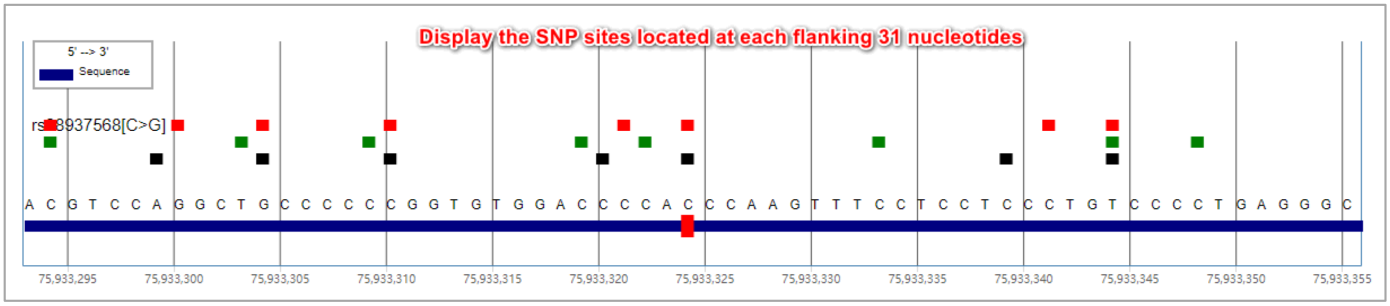


So far, **AsCRISPR** has integrated the genomes of Homo sapiens (hg19/GRCh37), Homo sapiens (hg38/GRCh38) and Mus musculus (mm10/GRCm38). **More genomes will be added for analysis in the near future.**

**Searching Criteria**

To identify guide sequences that may discriminate between two alleles, **AsCRISPR** explores both alleles and figures out if (i) mutations/SNPs give rise to novel PAMs, which confer stringent allele-specific targeting, and (ii) mutations/SNPs locate within the seed region of guide sequences, which may abolish the Cas cleavage. Users can freely set the length of seed sequence for each Cas type, to meet their personal need. The maximum lengths of seed sequences are set as follows:

- SpCas9, SaCas9 and their variants: 12 bp adjacent to a PAM
- AsCpf1, LbCpf1 and their variants: 6 bp adjacent to a PAM
- FnCpf1: 5 bp adjacent to a PAM
- Cas12b and its variants: 6 bp adjacent to a PAM
- CasX and its variants: 6 bp adjacent to a PAM

** Notably, previous studies have shown that **Cas12b and CasX exhibit greater specificity and possess broader seed sequence**, therefore the output number of allele-specific guide sequences under the criteria of 6 bp maximum seed sequence will actually be greatly underestimated.

#### Output of Guide Sequences

By filtering comparison results, **AsCRISPR** finally outputs candidate guide sequences that may target only one allele. The default output length of spacer sequences is set as 20-24 bp for different Cas9; 23 bp for AsCpf1/LbCpf1; 21 bp for FnCpf1; 23 bp for Cas12b; and 20 bp for CasX.

**AsCRISPR** ranks all guide sequences first by listing those with the same PAM back-to-back for better visualization. Alternatively, users can also freely re-rank their results according to their on-target efficiencies, off-targets or other properties, simply by clicking on the top of each category of the result table.

Se[illegible]GGTGTGGTACGCAGCC[C/A]CTTCGAGTACCCACAGTACTACCTGGCTG

Candidate guide sequences:
Mutations/SNPs are labeled in red and turned into lowercase
Can also re-rank guide sequences by targeting strand
Specificity Score
On-target efficiencies for SpCas9 or SaCas9
Potential off-targets (3 mismatches max.)
Restriction Enzymes

Exp[illegible]

| Ranking | Spacer Sequence | PAM | Cas9 Type | Targeting Strand | Direction | Gene Locus | GC content (%) | Specificity score | Xu 2015 score | Doench 2016 score | Moreno Mateos 2015 score | Azimuth in vitro score | Najm 2018 score | Self-complementarity | Off-targets 0-1-2-3 | Enzyme Information |
|---|---|---|---|---|---|---|---|---|---|---|---|---|---|---|---|---|
| 1 | ACGCAGCCcCTTCGA...ACC | CACAGTAC | CjCas9:NNNNRYAC | WT | + | exon:RHO | 65 | 1 | 0.37 | 0.53 | 0.76 | 0.57 | -- | 1 | 0-0-0-0 | Enzyme Information |
| 2 | ACGCAGCCaCTTCGAGTACC | CACAGTAC | CjCas9:NNNNRYAC | Varied | + | exon:RHO | 60 | 0.62 | 0.37 | 0.53 | 0.76 | 0.57 | -- | 1 | 0-1-0-0 | Enzyme Information |
| 3 | ACTGTGGGTACTCGAAGgGG | CTGCGTAC | CjCas9:NNNNRYAC | WT | - | exon:RHO | 60 | 1 | 0.8 | 0.69 | 0.87 | 0.7 | -- | 0 | 0-0-0-1 | Enzyme Information |
| 4 | ACTGTGGGTACTCGAAGtGG | CTGCGTAC | CjCas9:NNNNRYAC | Varied | - | exon:RHO | 55 | 0.84 | 0.8 | 0.69 | 0.87 | 0.7 | -- | 0 | 0-1-0-0 | Enzyme Information |
| 5 | GGGTACTCGAAGgGGCTGCG | TACCACAC | CjCas9:NNNNRYAC | WT | - | exon:RHO | 70 | 0.99 | 0.48 | 0.7 | 0.56 | 0.67 | -- | 0 | 0-0-0-1 | Enzyme Information |
| 6 | GGGTACTCGAAGtGGCTGCG | TACCACAC | CjCas9:NNNNRYAC | Varied | - | exon:RHO | 65 | 0.72 | 0.48 | 0.7 | 0.56 | 0.67 | -- | 0 | 0-1-0-0 | Enzyme Information |
| 7 | GGGTGTGGTACGCAGCCcCT | TCGAGTAC | CjCas9:NNNNRYAC | WT | + | exon:RHO | 70 | 0.99 | 0.54 | 0.51 | 0.77 | 0.7 | -- | 1 | 0-0-0-2 | Enzyme Information |
| 8 | GGGTGTGGTACGCAGCCaCT | TCGAGTAC | CjCas9:NNNNRYAC | Varied | + | exon:RHO | 65 | 0.84 | 0.54 | 0.51 | 0.77 | 0.7 | -- | 2 | 0-1-0-1 | Enzyme Information |
| 9 | ACGCAGCCcCTTCGA...ACC | CACAGT | SaCas9-KKH:NNNRRT | WT | + | exon:RHO | 65 | 1 | -- | -- | -- | -- | 0.71 | 1 | 0-0-0-5 | Enzyme Information |
| 10 | ACGCAGCCaCTTCGAGTACC | CACAGT | SaCas9-KKH:NNNRRT | Varied | + | exon:RHO | 60 | 0.95 | -- | -- | -- | -- | 0.71 | 1 | 0-1-0-1 | Enzyme Information |

Total 23 10/page < 1 2 3 >

To better display the differential points between the two alleles, in the result sheet, single-base mutations/SNPs are labeled in red and turned into lowercase, either located at the novel PAMs or spacer sequences. And for indel mutations, deleted nucleotides are replaced with asterisks (*) and labeled in red.

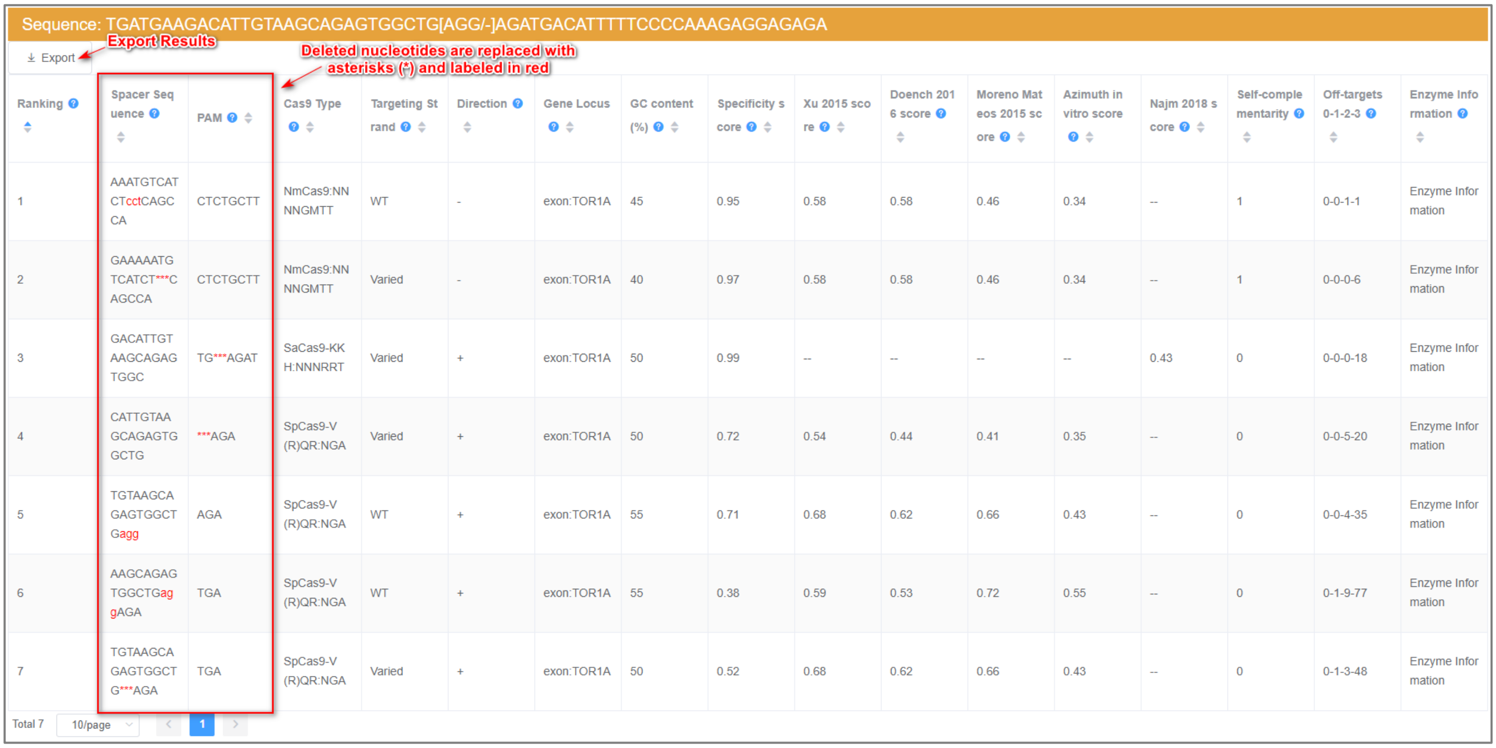


| Ranking | Spacer Sequence | PAM | Cas9 Type | Targeting Strand | Direction | Gene Locus | GC content (%) | Specificity score | Xu 2015 score | Doench 2016 score | Moreno Mateos 2015 score | Azimuth in vitro score | Najm 2018 score | Self-complementarity | Off-targets 0-1-2-3 | Enzyme Information |
|---|---|---|---|---|---|---|---|---|---|---|---|---|---|---|---|---|
| 1 | AAATGTCATCTcctCAGCCA | CTCTGCTT | NmCas9:NNNNGMTT | WT | - | exon:TOR1A | 45 | 0.95 | 0.58 | 0.58 | 0.46 | 0.34 | -- | 1 | 0-0-1-1 | Enzyme Information |
| 2 | GAAAAATGTCATCT***CAGCCA | CTCTGCTT | NmCas9:NNNNGMTT | Varied | - | exon:TOR1A | 40 | 0.97 | 0.58 | 0.58 | 0.46 | 0.34 | -- | 1 | 0-0-0-6 | Enzyme Information |
| 3 | GACATTGTAAGCAGAGTGGC | TG***AGAT | SaCas9-KKH:NNNRRT | Varied | + | exon:TOR1A | 50 | 0.99 | -- | -- | -- | -- | 0.43 | 0 | 0-0-0-18 | Enzyme Information |
| 4 | CATTGTAAGCAGAGTGGCTG | ***AGA | SpCas9-V(R)QR:NGA | Varied | + | exon:TOR1A | 50 | 0.72 | 0.54 | 0.44 | 0.41 | 0.35 | -- | 0 | 0-0-5-20 | Enzyme Information |
| 5 | TGTAAGCAGAGTGGCTGagg | AGA | SpCas9-V(R)QR:NGA | WT | + | exon:TOR1A | 55 | 0.71 | 0.68 | 0.62 | 0.66 | 0.43 | -- | 0 | 0-0-4-35 | Enzyme Information |
| 6 | AAGCAGAGTGGCTGaggAGA | TGA | SpCas9-V(R)QR:NGA | WT | + | exon:TOR1A | 55 | 0.38 | 0.59 | 0.53 | 0.72 | 0.55 | -- | 0 | 0-1-9-77 | Enzyme Information |
| 7 | TGTAAGCAGAGTGGCTG***AGA | TGA | SpCas9-V(R)QR:NGA | Varied | + | exon:TOR1A | 50 | 0.52 | 0.68 | 0.62 | 0.66 | 0.43 | -- | 0 | 0-1-3-48 | Enzyme Information |

Total 7 10/page 1

Notably, guide sequences will be reminded as “Not recommended” if (1) their GC contents are beyond 20%~80%; (2) containing four or more consecutive T, which might terminate the U6 or U3 promoter-drived transcription.

Sequence: AGGATCTGGTTCATTCTGTCTCACTTTTT[C/G]TTTAGGCTGCATTAGAAGTTAGCTTTCTT

Export

| Ranking | Spacer Sequence | PAM | Cas9 Type | Targeting Strand | Direction | Gene Locus | GC content (%) | Specificity score | Xu 2015 score | Doench 2016 score | Moreno Mateos 2015 score | Azimuth in vitro score | Najm 2018 score | Self-complementarity | Off-targets 0-1-2-3 | Enzyme Information |
|---|---|---|---|---|---|---|---|---|---|---|---|---|---|---|---|---|
| 11 | TGCAGCCTAAAcAAAAAGTG | AGA | SpCas9-V(R)QR:NGA | Varied | - | intron:TYR | 40 | 0.49 | 0.67 | 0.66 | 0.6 | 0.39 | -- | 2 | 0-1-3-55 | Enzyme Information |
| 12 | AATGCAGCCTAAAgAAAAAG | TGA | SpCas9-V(R)QR:NGA | WT | - | intron:TYR | 35 | 0.35 | 0.65 | 0.54 | 0.28 | 0.29 | -- | 2 | 0-1-6-84 | Enzyme Information |
| 13 | AATGCAGCCTAAAcAAAAAG | TGA | SpCas9-V(R)QR:NGA | Varied | - | intron:TYR | 35 | 0.6 | 0.65 | 0.54 | 0.28 | 0.29 | -- | 2 | 0-1-1-74 | Enzyme Information |
| 14 | TTCTGTCTCACTTTTTcTTT | AGG | SpCas9:NGG | WT | + | intron:TYR | 30 | 0.27 | 0.42 | 0.26 | 0.22 | 0.29 | -- | 0 | 0-1-49-290 | Enzyme Information |
| 15 | TTCTGTCTCACTTTTTgTTT | AGG | SpCas9:NGG | Varied | + | intron:TYR | 30 | 0.36 | 0.42 | 0.26 | 0.22 | 0.29 | -- | 0 | 0-1-25-202 | Enzyme Information |
| 16 | GCTAACTTCTAATGCAGCCT | AAAgAAA | StCas9:NNAGAAW | WT | - | intron:TYR | 45 | 0.98 | 0.53 | 0.5 | 0.17 | 0.3 | -- | 1 | 0-0-0-2 | Enzyme Information |
| 17 | CAGCCTAAAgAAAAAGTGAG | ACAGAAT | StCas9:NNAGAAW | WT | - | intron:TYR | 40 | 0.87 | 0.56 | 0.64 | 0.35 | 0.35 | -- | 2 | 0-0-1-13 | Enzyme Information |
| 18 | CAGCCTAAAcAAAAAGTGAG | ACAGAAT | StCas9:NNAGAAW | Varied | - | intron:TYR | 40 | 0.5 | 0.56 | 0.64 | 0.35 | 0.35 | -- | 2 | 0-1-1-7 | Enzyme Information |

Can check if more than 3 consecutive T existed in the guide, which might terminate the U6 or U3 promoter drived transcription.

Total 18 10/page < 1 2 >

dbSNP: GTCCAGGCTGCCCCCCGGTGTGGACCCCA[C/T]CCAAGTTTCCTCCTCCCTGTCCCCTGAGG

Export **dbSNP sequence #1 running results**

| Ranking | Spacer Sequence | PAM | Cas9 Type | Targeting Strand | Direction | Gene Locus | GC content (%) | Specificity score | Xu 2015 score | Doench 2016 score | Moreno Mateos 2015 score | Azimuth in vitro score | Najm 2018 score | Self-complementarity | Off-targets 0-1-2-3 | Enzyme Information |
|---|---|---|---|---|---|---|---|---|---|---|---|---|---|---|---|---|
| 1 | GAGGAGGAAACTTGGgTGGG | GTCCACAC | CjCas9:NNNNRYAC | Reference | - | exon:HSPB1 | 60 | 0.98 | 0.66 | 0.48 | 0.73 | 0.66 | -- | 0 | 0-0-0-6 | Enzyme Information |
| 2 | GAGGAGGAAACTTGGaTGGG | GTCCACAC | CjCas9:NNNNRYAC | Alternative | - | exon:HSPB1 | 55 | 0.84 | 0.66 | 0.48 | 0.73 | 0.66 | -- | 0 | 0-1-0-4 | Enzyme Information |
| 3 | CCCCCCGGTGTGGACCCCAc | CCAAGT | SaCas9-KKH:NNNRRT | Reference | + | exon:HSPB1 | 80 | 0.49 | -- | -- | -- | -- | 0.77 | 2 | 1-1-0-7 | Enzyme Information |
| 4 | CCCCCCGGTGTGGACCCCAt | CCAAGT | SaCas9-KKH:NNNRRT | Alternative | + | exon:HSPB1 | 75 | 0.88 | -- | -- | -- | -- | 0.77 | 1 | 0-2-1-1 | Enzyme Information |
| 5 | CAGGGAGGAGGAAACTTGGg | TGGGGT | SaCas9-KKH:NNNRRT | Reference | - | exon:HSPB1 | 60 | 0.84 | -- | -- | -- | -- | 0.81 | 0 | 0-0-3-45 | Enzyme Information |
| 6 | CAGGGAGGAGGAAACTTGGa | TGGGGT | SaCas9-KKH:NNNRRT | Alternative | - | exon:HSPB1 | 55 | 0.88 | -- | -- | -- | -- | 0.81 | 0 | 0-1-1-41 | Enzyme Information |
| 7 | CAGGGAGGAGGAAACTTGGg | TGGGGT | SaCas9:NNGRRT | Reference | - | exon:HSPB1 | 60 | 0.77 | -- | -- | -- | -- | 0.81 | 0 | 0-1-3-23 | Enzyme Information |
| 8 | CAGGGAGGAGGAAACTTGGa | TGGGGT | SaCas9:NNGRRT | Alternative | - | exon:HSPB1 | 55 | 0.84 | -- | -- | -- | -- | 0.81 | 0 | 0-1-3-11 | Enzyme Information |
| 9 | GGACAGGGAGGAGGAAACTT | GGa | SpCas9-V(R)QR:NGA | Alternative | - | exon:HSPB1 | 55 | 0.7 | 0.54 | 0.55 | 0.62 | 0.63 | -- | 1 | 0-0-3-38 | Enzyme Information |
| 10 | GGACAGGGAGGAGGAAACTT | GGg | SpCas9:NGG | Reference | - | exon:HSPB1 | 55 | 0.7 | 0.54 | 0.55 | 0.62 | 0.63 | -- | 1 | 0-0-8-74 | Enzyme Information |

**The recommended GC% is between 20%-80%, otherwise, it will be reminded.**

Total 15 10/page < 1 2 >

dbSNP: GTCCAGGCTGCCCCCCGGTGTGGACCCCA[C/G]CCAAGTTTCCTCCTCCCTGTCCCCTGAGG

Export **dbSNP sequence #2 running results**

| Ranking | Spacer Sequence | PAM | Cas9 Type | Targeting Strand | Direction | Gene Locus | GC content (%) | Specificity score | Xu 2015 score | Doench 2016 score | Moreno Mateos 2015 score | Azimuth in vitro score | Najm 2018 score | Self-complementarity | Off-targets 0-1-2-3 | Enzyme Information |
|---|---|---|---|---|---|---|---|---|---|---|---|---|---|---|---|---|
| 1 | GAGGAGGAAACTTGGgTGGG | GTCCACAC | CjCas9:NNNNRYAC | Reference | - | exon:HSPB1 | 60 | 0.98 | 0.66 | 0.48 | 0.73 | 0.66 | -- | 0 | 0-0-0-6 | Enzyme Information |
| 2 | GAGGAGGAAACTTGGcTGGG | GTCCACAC | CjCas9:NNNNRYAC | Alternative | - | exon:HSPB1 | 60 | 0.85 | 0.66 | 0.48 | 0.73 | 0.66 | -- | 0 | 0-1-0-1 | Enzyme Information |
| 3 | CCCCCCGGTGTGGACCCCAc | CCAAGT | SaCas9-KKH:NNNRRT | Reference | + | exon:HSPB1 | 80 | 0.49 | -- | -- | -- | -- | 0.77 | 2 | 1-1-0-7 | Enzyme Information |
| 4 | CCCCCCGGTGTGGACCCCAg | CCAAGT | SaCas9-KKH:NNNRRT | Alternative | + | exon:HSPB1 | 80 | 0.93 | -- | -- | -- | -- | 0.77 | 1 | 0-2-1-4 | Enzyme Information |
| 5 | CAGGGAGGAGGAAACTTGGg | TGGGGT | SaCas9-KKH:NNNRRT | Reference | - | exon:HSPB1 | 60 | 0.84 | -- | -- | -- | -- | 0.81 | 0 | 0-0-3-45 | Enzyme Information |
| 6 | CAGGGAGGAGGAAACTTGGc | TGGGGT | SaCas9-KKH:NNNRRT | Alternative | - | exon:HSPB1 | 60 | 0.92 | -- | -- | -- | -- | 0.81 | 0 | 0-1-2-27 | Enzyme Information |
| 7 | CAGGGAGGAGGAAACTTGGg | TGGGGT | SaCas9:NNGRRT | Reference | - | exon:HSPB1 | 60 | 0.77 | -- | -- | -- | -- | 0.81 | 0 | 0-1-3-23 | Enzyme Information |
| 8 | CAGGGAGGAGGAAACTTGGc | TGGGGT | SaCas9:NNGRRT | Alternative | - | exon:HSPB1 | 60 | 0.93 | -- | -- | -- | -- | 0.81 | 0 | 0-1-1-16 | Enzyme Information |
| 9 | GGACAGGGAGGAGGAAACTT | GGg | SpCas9:NGG | Reference | - | exon:HSPB1 | 55 | 0.7 | 0.54 | 0.55 | 0.62 | 0.63 | -- | 1 | 0-0-8-74 | Enzyme Information |
| 10 | AGGGAGGAGGAAACTTGGgT | GGG | SpCas9:NGG | Reference | - | exon:HSPB1 | 55 | 0.68 | 0.63 | 0.67 | 0.46 | 0.57 | -- | 0 | 0-1-6-62 | Enzyme Information |

**The recommended GC% is between 20%-80%, otherwise, it will be reminded.**

Total 15 10/page < 1 2 >

### On-target Efficiencies

The on-target efficiencies were calculated with multiple reported algorithms and were normalized to 0-1. For SpCas9, efficiency scores were predicted according to Xu et al., 2015; Doench et al., 2016; Moreno-Mateos et al., 2015; and Listgarden et al., 2018. For SaCas9, efficiency scores were predicted according to Najm et al., 2018. And for Cpf1, efficiency scores were predicted according to Kim et al., 2018. For Cas12b and CasX, efficiency scores were also predicted by DeepCpf1 according to Kim et al., 2018. **Note that Cas12b and CasX may have special efficiency scores which are different from those of Cas9 and Cpf1**, due to the lack of proved algorithms. Thus, those output DeepCpf1 scores will very likely not be applicable for them. We will definitely update the scoring system whenever there exists a convincing algorithm (or if you know one) to predict the on-target efficiencies of Cas12b and CasX.

**Nevertheless, the on-target efficiency scores provided here actually may only be used for on-target activity predictions.** If users have to choose certain guides for allele-specific targeting, they may choose and ignore those efficiency scores. However, if there are many specific guides for choice, users may look at and compare them with multiple properties, in which higher efficiency scores may be considered.

**** Notably, the analysis of query sequences that are not existed in any reference genome will not produce on-target efficiencies.**

### Off-targets and Specificity

The potential off-target sequences throughout the genome are searched by 3-base mismatches maximum. **AsCRISPR** lists the number of off-targets for each guide sequence with 0, 1, 2 or 3 mismatches (0-1-2-3). Clicking on the (0-1-2-3) will reveal more information about the off-target information, including the locations (exon, intron or intergenic region), sequence mismatches and so forth.

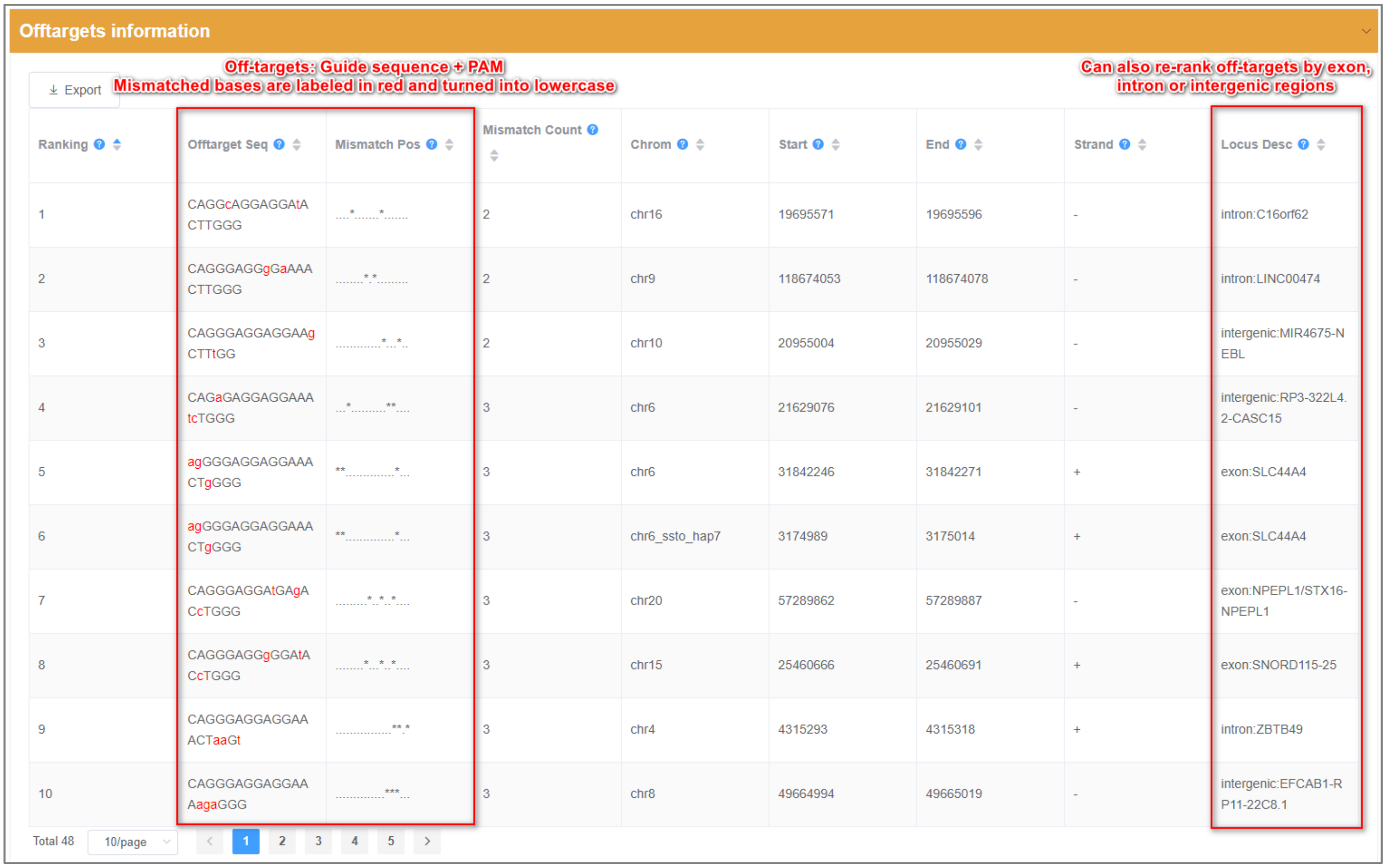


| Ranking | Offtarget Seq | Mismatch Pos | Mismatch Count | Chrom | Start | End | Strand | Locus Desc |
|---|---|---|---|---|---|---|---|---|
| 1 | CAGGcAGGAGGAtACTTGGG | ....*.......*....... | 2 | chr16 | 19695571 | 19695596 | - | intron:C16orf62 |
| 2 | CAGGGAGGgGaAAACTTGGG | ........*.*......... | 2 | chr9 | 118674053 | 118674078 | - | intron:LINC00474 |
| 3 | CAGGGAGGAGGAAgCTTtGG | .............*...*.. | 2 | chr10 | 20955004 | 20955029 | - | intergenic:MIR4675-NEBL |
| 4 | CAGaGAGGAGGAAAtcTGGG | ...*..........**.... | 3 | chr6 | 21629076 | 21629101 | - | intergenic:RP3-322L4.2-CASC15 |
| 5 | agGGGAGGAGGAAACTgGGG | **.............*... | 3 | chr6 | 31842246 | 31842271 | + | exon:SLC44A4 |
| 6 | agGGGAGGAGGAAACTgGGG | **.............*... | 3 | chr6_ssto_hap7 | 3174989 | 3175014 | + | exon:SLC44A4 |
| 7 | CAGGGAGGAtGAgACcTGGG | .........*..*..*.... | 3 | chr20 | 57289862 | 57289887 | - | exon:NPEPL1/STX16-NPEPL1 |
| 8 | CAGGGAGGgGGAtACcTGGG | ........*...*..*.... | 3 | chr15 | 25460666 | 25460691 | + | exon:SNORD115-25 |
| 9 | CAGGGAGGAGGAAACTaaGt | ................**.* | 3 | chr4 | 4315293 | 4315318 | + | intron:ZBTB49 |
| 10 | CAGGGAGGAGGAAAagaGGG | ..............***... | 3 | chr8 | 49664994 | 49665019 | - | intergenic:EFCAB1-RP11-22C8.1 |

Total 48 10/page < 1 2 3 4 5 >

The specificity score measures the uniqueness of a guide sequence in the genome. The higher the specificity score, the lower are off-target effects. Specificity scores were calculated based on Hsu scores by Hsu et al., 2013 and CFD scores by Donech et al., 2016. For Cpf1, Cas12b and CasX, no off-target ranking algorithms were available in the literatures so far, instead we just applied Hsu and CFD scores to their off-targets. We will update the scoring system whenever there exists a convincing algorithm (or if you know one) to predict the off-target effects for Cpf1, Cas12b and CasX.

Sequence: CTCCAATGCGACGGGTGTGGTACGCAGCC[C/A]CTTCGAGTACCCACAGTACTACCTGGCTG

Export

| Ranking | Spacer Sequence | PAM | Cas9 Type | Targeting Strand | Direction | Gene Locus | GC content (%) | Specificity score | Xu 2015 score | Doench 2016 score | Moreno Mateos 2015 score | Azimuth in vitro score | Najm 2018 score | Self-complementarity | Off-targets 0-1-2-3 | Enzyme Information |
|---|---|---|---|---|---|---|---|---|---|---|---|---|---|---|---|---|
| 11 | CAGGTAGTACTGTGGGTACT | CGAAGt | SaCas9-KKH:NNNRRT | Varied | - | exon:RHO | 50 | 0.94 | -- | -- | -- | -- | 0.38 | 1 | 0-0-1-16 | Enzyme Information |
| 12 | GGGTGTGGTACGCAGCCcCT | TCGAGT | SaCas9-KKH:NNNRRT | WT | + | exon:RHO | 70 | 1 | -- | -- | -- | -- | [illegible] | 1 | 0-0-0-3 | Enzyme Information |
| 13 | GGGTGTGGTACGCAGCCaCT | TCGAGT | SaCas9-KKH:NNNRRT | Varied | + | exon:RHO | 65 | 0.97 | -- | -- | -- | -- | 0.62 | 2 | 0-1-1-4 | Enzyme Information |
| 14 | GGGTGTGGTACGCAGCCcCT | TCGAGT | SaCas9:NNGRRT | WT | + | exon:RHO | 70 | 0.95 | -- | -- | -- | -- | 0.62 | 1 | 0-0-2-2 | Enzyme Information |
| 15 | GGGTGTGGTACGCAGCCaCT | TCGAGT | SaCas9:NNGRRT | Varied | + | exon:RHO | 65 | 0.97 | -- | -- | -- | -- | 0.62 | 2 | 0-1-0-1 | Enzyme Information |
| 16 | GGTGTGGTACGCAGCCcCTT | CGAG | SpCas9-EQR:NGAG | WT | + | exon:RHO | 65 | 1 | 0.45 | 0.44 | 0.66 | 0.61 | -- | 1 | 0-0-0-0 | Enzyme Information |
| 17 | GGTGTGGTACGCAGCCaCTT | CGAG | SpCas9-EQR:NGAG | Varied | + | exon:RHO | 60 | 0.72 | 0.45 | 0.44 | 0.66 | 0.61 | -- | 2 | 0-1-0-1 | Enzyme Information |
| 18 | GGTGTGGTACGCAGCCcCTT | CGA | SpCas9-V(R)QR:NGA | WT | + | exon:RHO | 65 | 0.93 | 0.45 | 0.44 | 0.66 | 0.61 | -- | 1 | 0-0-0-6 | Enzyme Information |
| 19 | GGTGTGGTACGCAGCCaCTT | CGA | SpCas9-V(R)QR:NGA | Varied | + | exon:RHO | 60 | 0.69 | 0.45 | 0.44 | 0.66 | 0.61 | -- | 2 | 0-1-0-5 | Enzyme Information |
| 20 | CTGTGGGTACTCGAAGgGGC | TGCG | SpCas9-VRER:NGCG | WT | - | exon:RHO | 65 | 1 | 0.59 | 0.47 | 0.46 | 0.29 | -- | 0 | 0-0-0-0 | Enzyme Information |

Total 23 | 10/page | 1 2 3

Compromised Specificity Score

1-base mismatch
Read WT sequence as an off-target

**** Note: When processing with the guide sequences from varied/alternative alleles, it is highly possible that one of those off-targets (with only 1-base mismatch) would be exactly the sequence of their corresponding WT/reference alleles. This will compromise the specificity scores of guide sequences from varied/alternative alleles to some extent (see the above Figure).**

#### Restriction Enzyme Sites

**AsCRISPR** displays the restriction enzyme sites along the spacer sequences, which might be disrupted after gene targeting, and further determine whether those candidate enzymes are specifically targeted either allele. This provides an important tool for RFLP-based characterization and screening for targeted single colonies.

**Enzymes information**

Export

Totally 8 commercial suppliers for choice

Can determine whether candidate restriction enzymes are specifically targeted to this allele

| Ranking | Enzyme name | Recognition sequence | Supplier | Incubation Temperature | Recommended Buffer | Allele Specific |
|---|---|---|---|---|---|---|
| 1 | AvaII | GGWCC | NEB,Progema,Roche,Thermo,TaKaRa,Sigma | 37,37,37,37,37,37 | CutSmart,C,A,F,M,SA | No |
| 2 | BmgT120I | GGNCC | TaKaRa | 37 | H | No |
| 3 | BseRI | GAGGAG | NEB | 37 | CutSmart | No |
| 4 | BspLI | GGNNCC | Thermo | 37 | T | No |
| 5 | Cfr13I | GGNCC | ToYoBo,Thermo | 37,37 | TA + BSA,T | No |
| 6 | CviKI-1 | RGCY | NEB | 37 | CutSmart | Yes; Alternative Strand Specific |
| 7 | Eco47I | GGWCC | Thermo,ToYoBo | 37,37 | R,H | No |
| 8 | Hpy166II | GTNNAC | NEB | 37 | CutSmart | No |
| 9 | Hpy8I | GTNNAC | Thermo | 37 | T | No |
| 10 | NlaIV | GGNNCC | NEB,Thermo | 37,37 | CutSmart,F | No |

Total 13 10/page 1 2

#### Dominant Database

**AsCRISPR** collected dominant SNV sites from ClinVar and OMIM databases for allele-specific analysis, and generated a ***Dominant Database*** depositing candidate targetable sgRNAs, which may specifically target the alternative allele for each dominant SNV site. Users can search by Disease name, Gene name or rsID for targetable site.

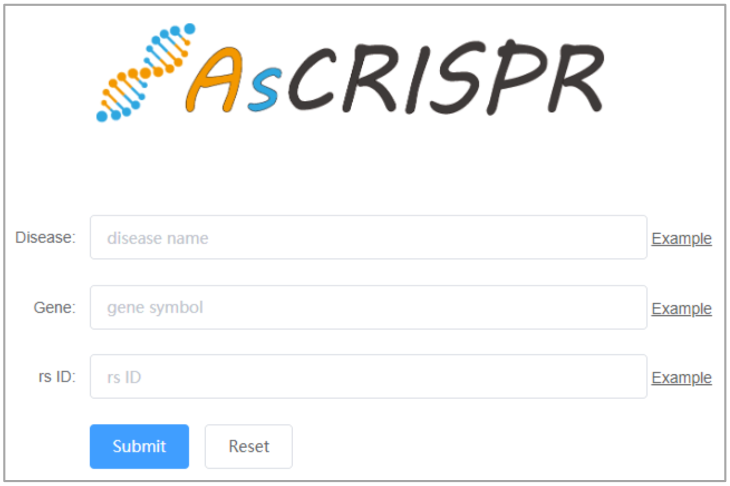
